# Persistent confined migration confers permanent nuclear and functional changes in migrating cells

**DOI:** 10.1101/2023.01.05.522838

**Authors:** Ana de Lope-Planelles, Raquel González-Novo, Elena Madrazo, Héctor Zamora-Carreras, Verónica Torrano, Horacio López-Menéndez, Pedro Roda-Navarro, Francisco Monroy, Javier Redondo-Muñoz

**Affiliations:** Department of Molecular Medicine, Centro de Investigaciones Biológicas Margarita Salas (CIB Margarita Salas-CSIC), Madrid, Spain; Department of Immunology, School of Medicine, University Complutense de Madrid and 12 de Octubre Health Research Institute (Imas12) Madrid, Spain; Department of Biochemistry and Molecular Biology, University of the Basque Country, Leioa, Spain; Department of Physical Chemistry, Complutense University, Madrid, Spain; Translational Biophysics, Hospital Doce de Octubre Health Research Institute (imas12), Madrid, Spain

**Author notes:** **Corresponding author**: Javier Redondo-Muñoz.

**Keywords:** Cell migration, nuclear deformability, mechanobiology, lamin, DNA damage.

## Abstract

Nuclear deformability plays a critical role in cell migration. During this process, the remodeling of internal components of the nucleus has a direct impact on DNA damage and cell behavior; however, how persistent migration promotes nuclear changes leading to phenotypical and functional consequences remains poorly understood. Here, we described that the persistent migration through physical barriers was sufficient to promote permanent modifications in migratory-altered cells. We found that lamin B1 altered its localization, concomitant with morphological and transcriptional changes. Migratory-altered cells showed alterations in cellular functions such as DNA repair and cell migration. We applied biochemical and biophysical approaches to identify that confined conditions altered the biomechanical response of the nucleus. Mechanistically, we determined that actin dynamics controlled the redistribution of lamin, and the basal levels of DNA damage in migratory-altered cells. Our observations reveal a novel role for confined cell conditions in consistent nuclear and genomic alterations that might handle the genetic instability and cellular heterogeneity in aging diseases and cancer.

**Highlights:**

- Persistent confined migration promotes permanent mophological changes.
- Lamin B1 is redistribution in the nucleus of migratory altered cells.
- Migratory-altered cells exhibit transcriptional and functional changes related to cell migration and survival.
- Actin polymerization controls nuclear changes induced by cell migration.

## Introduction

Cell migration is a fundamental process involved in physiological and pathological conditions such as development, immune response, inflammation, and cancer invasion [1, 2]. Migrating cells have to cross through different tissue environments, including the extracellular matrix and endothelial barriers, which provide multiple biochemical and mechanobiological signals for an active and effective cell migration [3, 4]. Nuclear deformability has been defined as a major limiting factor for cell migration across physical barriers [5]. The nucleus is a highly dynamic organelle composed of the nuclear envelope, the lamina network, and the chromatin structure [6–8]. The cytoskeleton exerts and transmits mechanical forces and external stimuli into the nucleus [9, 10]; and previous studies have demonstrated that nuclear deformation induced by mechanical stress regulates chromatin changes, DNA damage response, and cell stemness [11–15]. Likely, several human pathologies are associated with defects in the response of the nucleus to mechanical stimuli, including progeria, cardiomyopathies, and cancer [16, 17]. However, how much persistent migration might control more stable nuclear changes remains poorly understood.

On this basis, we compare two cell migration settings, one-round migration (ORM) cells, and migratory-altered (MA) cells, which might promote nuclear changes while regulating the functional and mechanical responses of the cell in two differential mechanobiological scenarios. We have found that cell migration triggers a change in the distribution of lamin B1 and the accumulation of DNA damage markers. To elucidate the functional consequences of nuclear changes induced by persistent cell migration, we explored how moving cells might alter their gene expression signature and other cellular functions. We demonstrated that the mechanical stress during cell migration generates permanent changes in the transcriptional profile of migratory-altered cells. Furthermore, strong nuclear deformability under persistent cell migration alters a range of mechanical and functional responses, including cell migration, and DNA damage repair. Noticeably, latrunculin treatment causes lamin B1 localization at the nuclear periphery and rescues the basal phenotype and morphology of the nucleus and chromatin compaction, suggesting that actin polymerization is critical for some of these processes. Overall, our experimental findings offer insight into how nuclear deformability induced by cell migration contributes to genomic alterations, which could lead to functional changes, such as tumor progression and heterogeneity of cancer cells.

## Results

### Mechanical constriction promotes morphological changes in the nucleus of migrating cells

We first determined how confined migration alters the nuclear morphology of migrating cells. We seeded leukemia cells on rigid filter membranes and allowed their migration (one-round migration, ORM) through the pores for 24 h. Then, we collected these cells and compared them with their non-migrating counterparts to further characterize how cell migration through physical barriers might influence the nucleus of leukemia cells. First, we observed that ORM cells increased their nuclei in comparison to non-migrating cells (Figures 1A, 1B, S1A and S1B). It has been shown that changes in the size of isolated nuclei can be determined by flow cytometry [18], and we verified that isolated nuclei from ORM cells showed bigger size than nuclei from non-migrating cells (Figure S1C). To understand the long-term consequences of persistent cell migration, and whether a consistent migratory stimulus might induce similar changes to ORM cells, we generated stable migratory-altered (MA) cells, by allowing cells to migrate three–rounds across rigid filter membranes and then culturing them in the absence of further mechanical stimuli (Figure 1C). We visualized the nucleus of MA cells and confirmed that persistent cell migration also induced an increment in the size of the nucleus (Figures 1D, 1E, S1C, and S1D). These results were in line with our observations above, which showed that ORM cells presented bigger nuclei compared to control cells. Together, these data indicate that cell squeezing during migration induces changes in the nuclear size of moving cells. Having observed differences in the size of the nuclei, we next focused in the nuclear morphology of MA cells. Electron microscopy images revealed a remarkable aberrant morphology of the nucleus of MA cells compared to those from control cells (Figures 1F, and S1E); although we did not observe differences in the nuclear morphology of ORM cells (Figures S1F, and S1G). This suggests that increasing nuclear size is a common effect of confined migration, whilst altering nuclear morphology might only be characteristic in the context of persistent migration.

**Figure 1:**
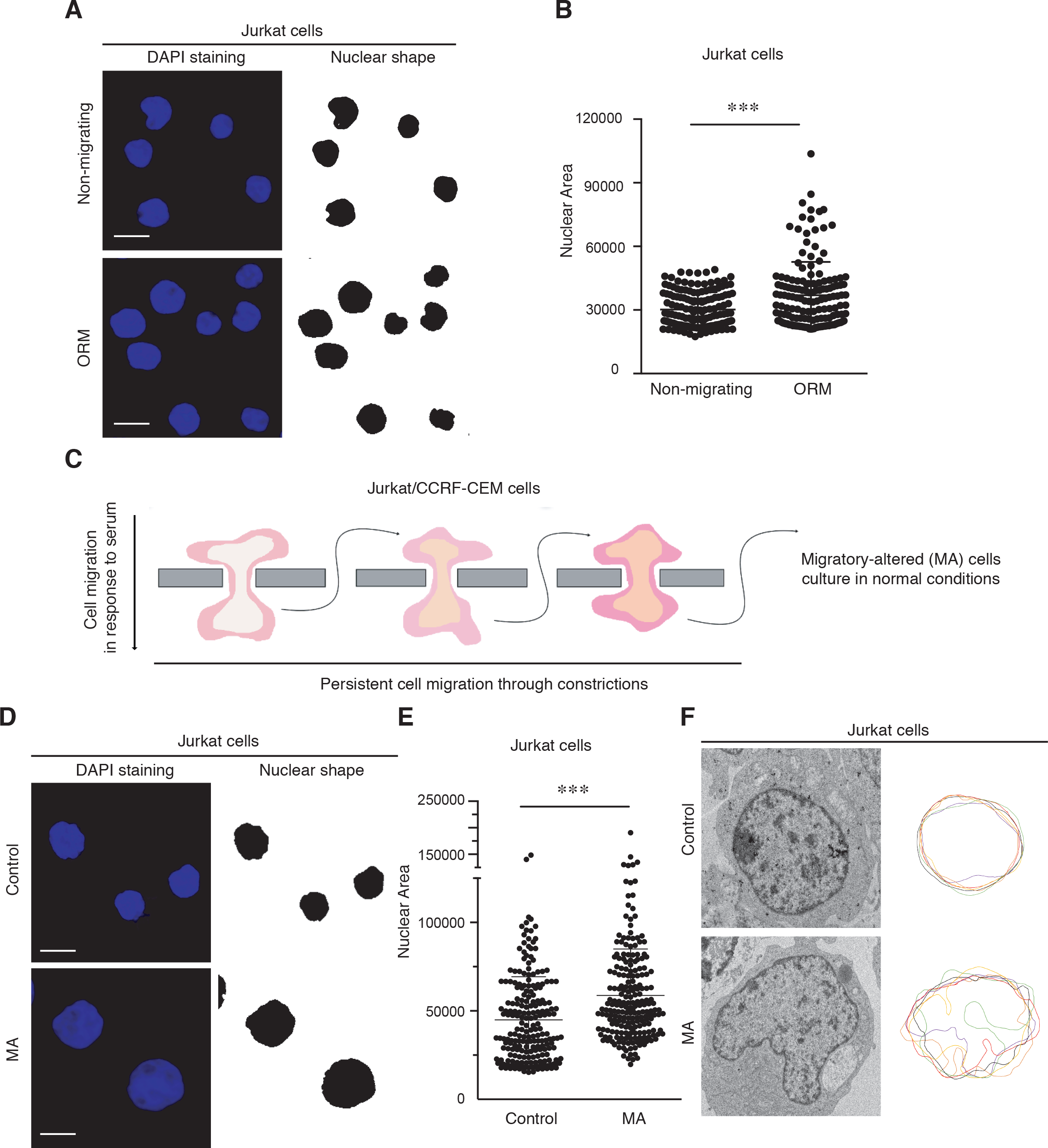
Cell migration through constrictions governs nuclear changes in leukemia cells. **(A)** Jurkat cells were allowed to migrate across 3 μm transwell inserts for 24h. Non-migrating and one-round migrated (ORM) cells were collected from the upper and bottom chambers, respectively, sedimented on poly-L-lysine coated coverslips, fixed and stained with DAPI (blue) for their analysis by confocal microscopy. Right panels indicate in black the area of the nuclei. Bar 10 μm. **(B)** Graph shows changes in the nuclear area of Jurkat cells upon one-round of migration through constrictions. Mean n= 157 cells ± SD (3 replicates). **(C)** Jurkat and CCRF-CEM cells were forced to migrate several rounds through 3 μm transwell inserts. Then, migrated cells were collected as migratory-altered (MA) cells, expanded and kept in culture conditions. **(D)** Control and MA Jurkat cells were sedimented on poly-L-lysine coated coverslips, fixed and stained with DAPI. Right panels indicate in black the area of the nuclei. Bar 10 μm. **(E)** Graph shows changes in the nuclear area of control and MA cells. Mean n= 216 cells ± SD (5 replicates). **(F)** Control and MA Jurkat cells were collected and processed for thin section electron microscopy to visualize the nuclear morphology. Plot shows changes in the nuclear circularity of control and MA Jurkat cells (n=6 representative cells).

### Persistent migration alters permanently the lamin B1 distribution within the nucleus

Since nuclear lamina controls the mechanical properties and behavior of the nucleus [19], we next sought to elucidate whether confined migration promoted alterations in the lamina network and found that constricted migration altered the distribution of lamin B1 in ORM cells (Figures 2A, and S2A). While non-migrating cells showed two characteristic peaks for lamin B1 signal in the transversal section of the nucleus (corresponding to the nuclear periphery), ORM cells presented an aberrant distribution of lamin B1 (Figures 2B, S2B, and Movies S1, S2). Temporal changes in internal components might critically regulate the morphology of the nucleus in moving cells; therefore, the nuclear volume and lamin B1 redistribution observed would be recovered in ORM cells once the migratory effect disappears. We demonstrated that ORM cells cultured in suspension for an additional 24 hours reduced their nuclear size to levels comparable to those from non-migrating cells (Figures 2C, 2D, S2C, and S2D). Furthermore, these cells also recovered the normal localization of lamin B1 at the nuclear periphery (Figures 2E, and S2E). To investigate the effect of persistent migration on the nuclear changes of MA cells, we determined their lamin B1 distribution and found that MA cells showed an aberrant distribution of lamin B1 (Figures 2F, 2G, S2F, S2G, and Movies S3, S4). These findings led us to conclude that persistent migration is a critical factor to induce permanent alterations in moving cells, whilst transient migration shows only a temporary effect on the nucleus of migrating cells. Altogether, these results indicate that moving cells derived both from transient and persistent migration underwent lamin B1 redistribution, which can affect multiple cellular functions.

**Figure 2:**
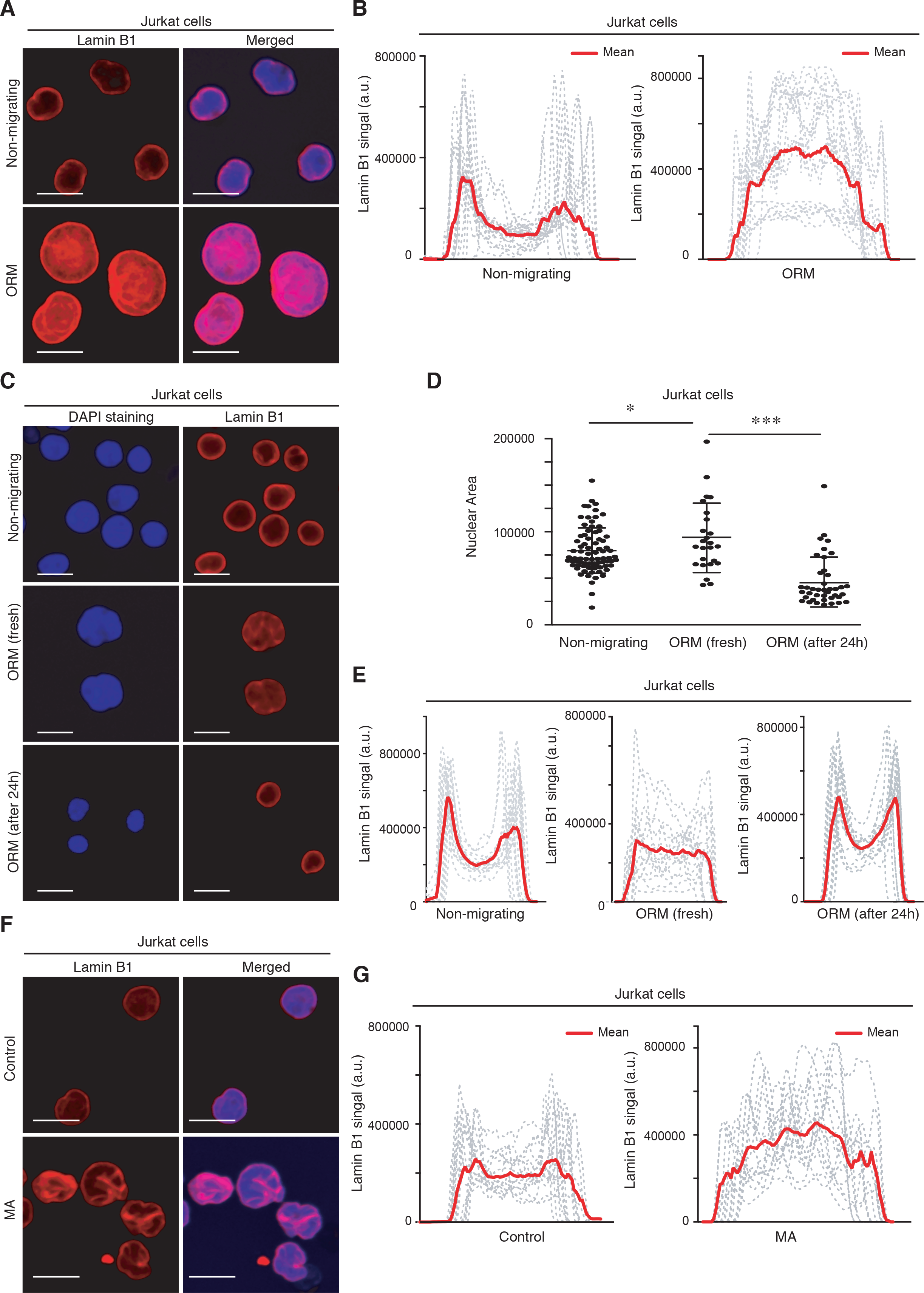
Confined migration alters the lamin B1 distribution of moving cells. (A) Non-migrating and ORM Jurkat cells were seeded on poly-L-lysine-coated glasses and stained with DAPI (blue) and anti-lamin B1 antibody (red) for their analysis by confocal microscopy. Bar 10 μm. (B) Line plots show the signal profile of lamin B1 from 15 representative nuclei. Red line indicates the mean intensity of the profiles analyzed. (C) Control, fresh ORM, and ORM Jurkat cells collected and cultured in suspension for an additional 24 h were seeded on poly-L-lysine-coated glasses and analyzed by confocal microscopy. Bar 10 μm. (D) Graph shows changes in the nuclear area of the cells from (C). Mean n=25-92 cells ± SD (2 independent replicates). (E) Line plots show the signal profile of lamin B1 from 15 representative nuclei of the cells from (C). (F) Control and MA Jurkat cells were seeded on poly-L-lysine-coated glasses and analyzed by confocal microscopy. Bar 10 μm. (G) Line plots show the signal profile of lamin B1 from 15 representative nuclei of control and MA Jurkat cells. The red line indicates the mean intensity of the profiles analyzed.

### Persistent migration induces enduring changes in the transcriptional profile of migrating cells

It has been previously described that nuclear shape regulates the transcriptional activity of cells [20, 21]. As MA cells showed a different and sustained phenotype compared to control cells, we hypothesized that persistent cell migration might produce enduring changes in the transcriptional landscape of MA cells. We evaluated the expression levels of transcripts of control or MA cells and found a significant differential expression of 1.8-fold change in 639 genes (285 up- and 354 down-regulated) in MA cells compared with control cells (Figures 3A, S3A, and Table S1). We next extended our observation to KEGG (Kyoto Encyclopedia of Genes and Genomes) and GO (Gene Ontology) enrichment analyses, which revealed that most of the altered genes were connected to cell division, survival, DNA damage checkpoints and protein modification (Figure S3B). To confirm changes in gene expression identified by microarray analysis, we measured the relative mRNA levels of two genes related to leukemia (*EZH2* and *JAK2*), as well as the expression levels of 5 randomly selected proteins (Figures 3C, S3C, and S3D). Once validated these observations, our pathway analysis demonstrated that upregulated genes of MA cells encode proteins related to G2/M progression, mitotic spindle, and chromosome segregation (Figure 3B, and Table S2). Previous reports showed that confined conditions regulate cell proliferation and cell cycle progression [22, 23]; however, we observed no differences in the proliferative properties of MA cells compared to control cells (Figures 3D, 3E, S3E, and S3F). We validated that the cell cycle progression was not altered in MA cells (Figures 3F, and S3G), suggesting that cell proliferation might be affected by other external stimuli, rather than just potential transcriptional changes. Overall, our results indicate that persistent cell migration might induce changes in the transcriptional profile of moving cells.

**Figure 3:**
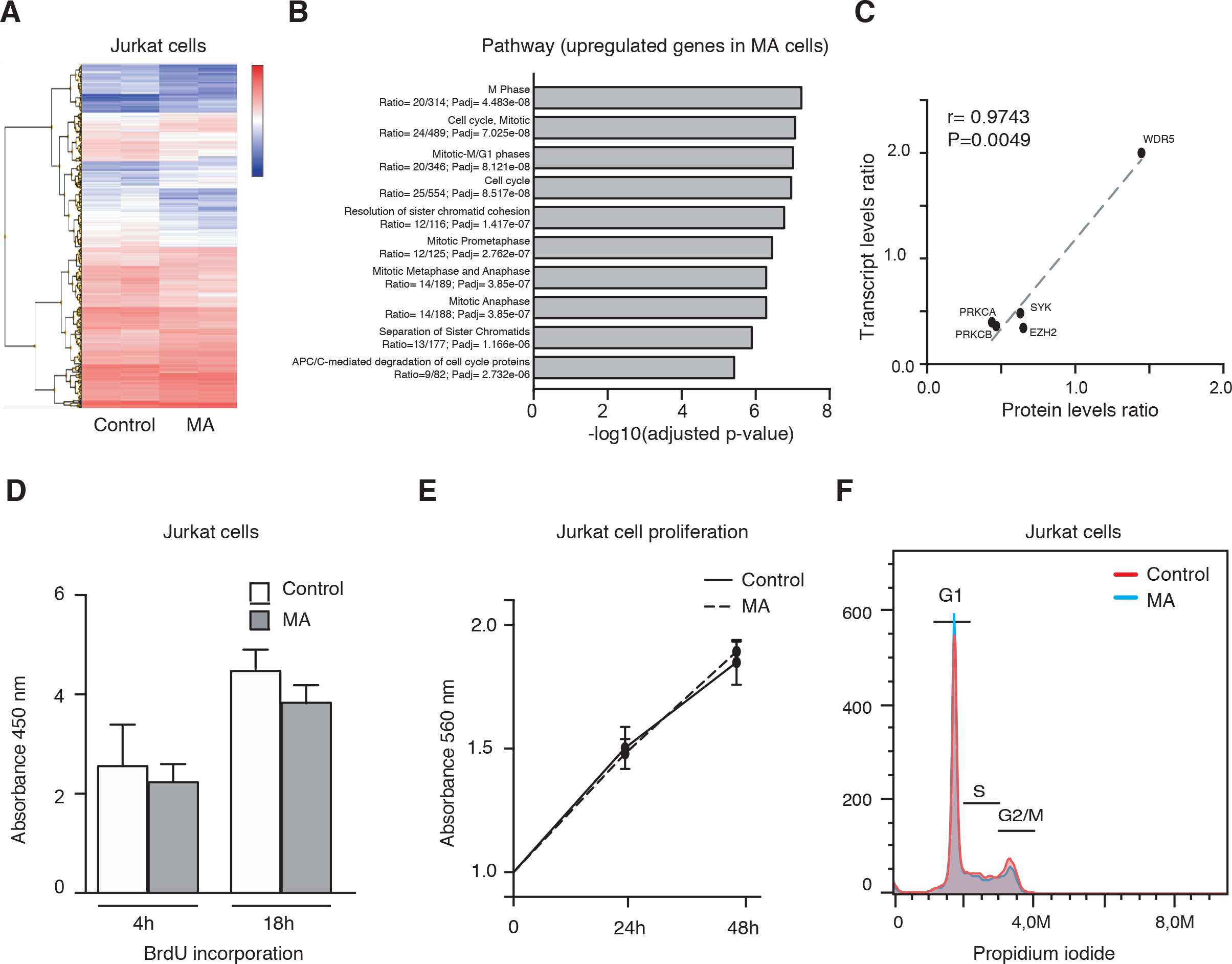
Persistent migration induces enduring changes in the transcriptional profile of migrating cells. **(A)** Control and MA Jurkat cells were lysed, and transcriptional changes determined by mRNA expression microarray. Heat map shows the relative gene expression patterns of control and MA cells. **(B)** Pathway enrichment analysis based on differentially upregulated genes in MA Jurkat cells. **(C)** Validation of microarray data by protein expression levels. Plot shows the transcript-protein level pairs from 5 randomly selected proteins from the transcriptional analysis by microarray. Gene names, Pearson correlation analysis (r) and P value are indicated. **(D)** Control and MA Jurkat cells were incubated with BrdU for 4 and 18 h. Then, cells were fixed and BrdU incorporation quantified. Mean n = 6 replicates ± SD. **(E)** Control (dark line) and MA (dashed line) Jurkat cells were cultured at indicated times and cell proliferation was quantified by MTT assay. Mean n = 3 replicates ± SD. **(F)** Control (in red) and MA (in blue) Jurkat cells were fixed, permeabilized and stained with propidium iodide. Then, cell cycle progression was analyzed by flow cytometry. Graph shows the G1, S and G2/M phases according to DNA content.

### Migrating cells show altered basal levels of DNA damage markers and resistance to apoptosis

In addition to controlling the cell cycle progression, cell migration is also known to favor DNA damage upon extreme nuclear deformability [24]. We investigated whether cell migration might increase the constitutive levels of γH2AX (a well-known DNA damage marker) and found that most of the non-migrating cells showed low levels of γH2AX (<2 foci per nucleus), in contrast to fresh ORM cells (Figures 4A). Likely, the percentage of positive MA cells for γH2AX was higher than control cells (Figures 4B, and 4C), indicating that cell migration increases the basal levels of DNA damage in moving cells. This suggests that cells undergoing transient and persistent migration increase their basal levels of DNA damage, which would presumably have an impact on how leukemia cells respond to therapy. Our pathway analysis showed that MA cells might downregulate genes involved in endoplasmic reticulum stress and unfolded protein response (Figures 4D and Table S2), which might be relevant for drug resistance in cancer (25). To gain insight into how persistent migration might affect cell viability, we treated control and MA cells with methotrexate, a DNA biosynthesis inhibitor used in clinics against multiple human pathologies [26]. Interestingly, we found that MA cells showed higher resistance to apoptosis than control cells (Fig. 4E, 4F). While this is a simple model to induce apoptosis, we could speculate that MA cells would turn less sensitive to some conventional therapies compared to control cells. Overall, these observations indicate that persistent migration would facilitate the resistance to apoptosis and the adaptation to conventional therapies of invasive moving cells.

**Figure 4:**
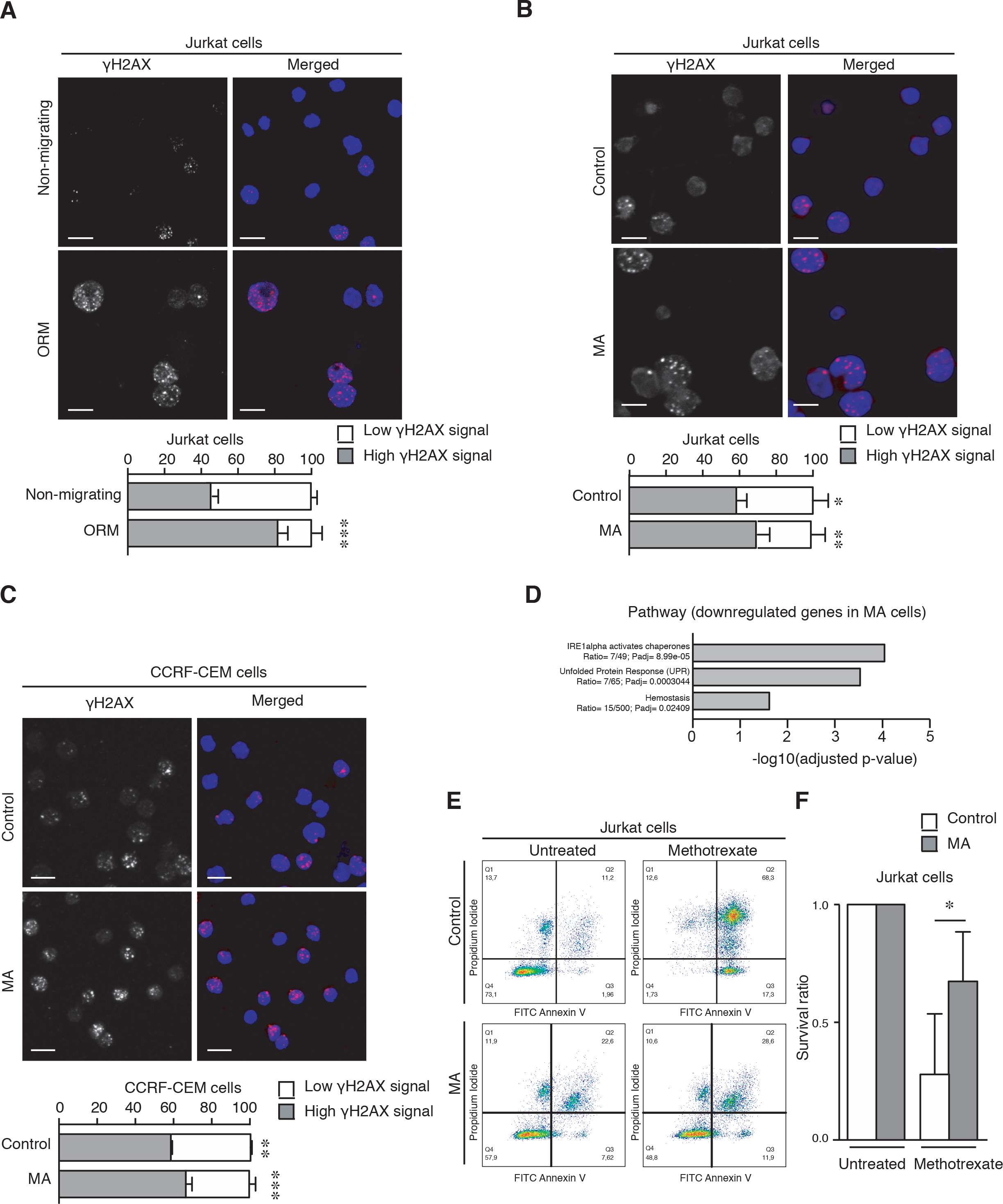
Persistent cell migration leads to aberrant DNA damage response. **(A)** Non-migrating and ORM Jurkat cells were seeded on poly-L-lysine-coated glasses and stained with DAPI (blue) and anti-γH2AX antibody (red) for their analysis by confocal microscopy. Bar 10 μm. Graph shows the percentage of cells with more than 2 visible foci for γH2AX (dark grey, as high γH2AX signal). Mean n = 116-118 ± SD (3 replicates). **(B)** Control and MA Jurkat cells were seeded on poly-L-lysine-coated glasses, stained, and analyzed by confocal microscopy. Bar 10 μm. Graph shows the percentage of control and MA Jurkat cells with more than 2 visible foci for γH2AX. Mean n = 28-35 cells ± SD (2 replicates). **(C)** Control and MA CCRF-CEM cells were seeded on poly-L-lysine-coated glasses glasses, stained, and analyzed by confocal microscopy. Bar 10 μm. Graph shows the percentage of control and MA CCRF-CEM cells with more than 2 visible foci for γH2AX. Mean n = 146-171 cells ± SD (3 replicates). **(D)** Pathway enrichment analysis based on differentially downregulated genes in MA Jurkat cells. **(E)** Control and MA Jurkat cells were cultured in the presence or not of methotrexate (1 μM) for 24 h. Then, cells were collected and stained with annexin V-FITC and propidium iodide for their flow cytometry analysis. **(F)** Graph shows the survival ratio of control and MA Jurkat cells upon normalization to their untreated conditions. Mean n = 3 ± SD.

### Migrating cells show different biomechanical responses of the nucleus and altered cell invasiveness

Lamins and chromatin contribute to the mechanical properties and size of the nucleus [27], and we hypothesized that confined migration might control not only the nuclear morphology but also have a direct impact on the mechanical behavior of the nucleus. We assayed an external mechanical pressure technology to exert forces onto isolated nuclei from control and MA cells [28], and observed that the nuclei of MA cells increased 2.5-fold under external pressure, whilst control cells showed only 1.77-fold (Figures 5A, 5B, S4A, and S4B). This indicates that MA cells showed a higher nuclear deformation capacity compared to control cells. Then, we trapped isolated nuclei from control and MA cells and analyzed them with optical tweezers. Unexpectedly, isolated nuclei from MA cells showed no differences in their stiffness and water diffusivity compared to control cells (Figures 5C, and 5D). It is possible that the response generated by optical tweezers represented a different mechanical behavior than the whole compression of the nucleus. To further characterize the mechanical behavior of the nucleus, we addressed its response under osmotic stress. The balance of cations regulates the osmotic pressure and the chromatin compaction in the nucleus [29]. We observed that isolated nuclei from MA cells responded better to the addition of cations (42% of shrinking compared to 10% of nuclei from control cells), whilst the addition of EDTA promoted higher swelling in isolated nuclei from control cells rather than from MA cells (60% and 30% of increment in the nuclear area, respectively; Figures 5E, and 5F). To compare these findings with transitory changes induced by cell migration, we isolated nuclei from ORM cells and observed an increment in their nuclear deformability area upon compression by external forces (Figures S4C, and S4D). Furthermore, isolated nuclei from ORM cells showed higher nuclear stiffness and diminished water diffusivity compared to non-migrating cells (Figures S4E, and S4F).

**Figure 5:**
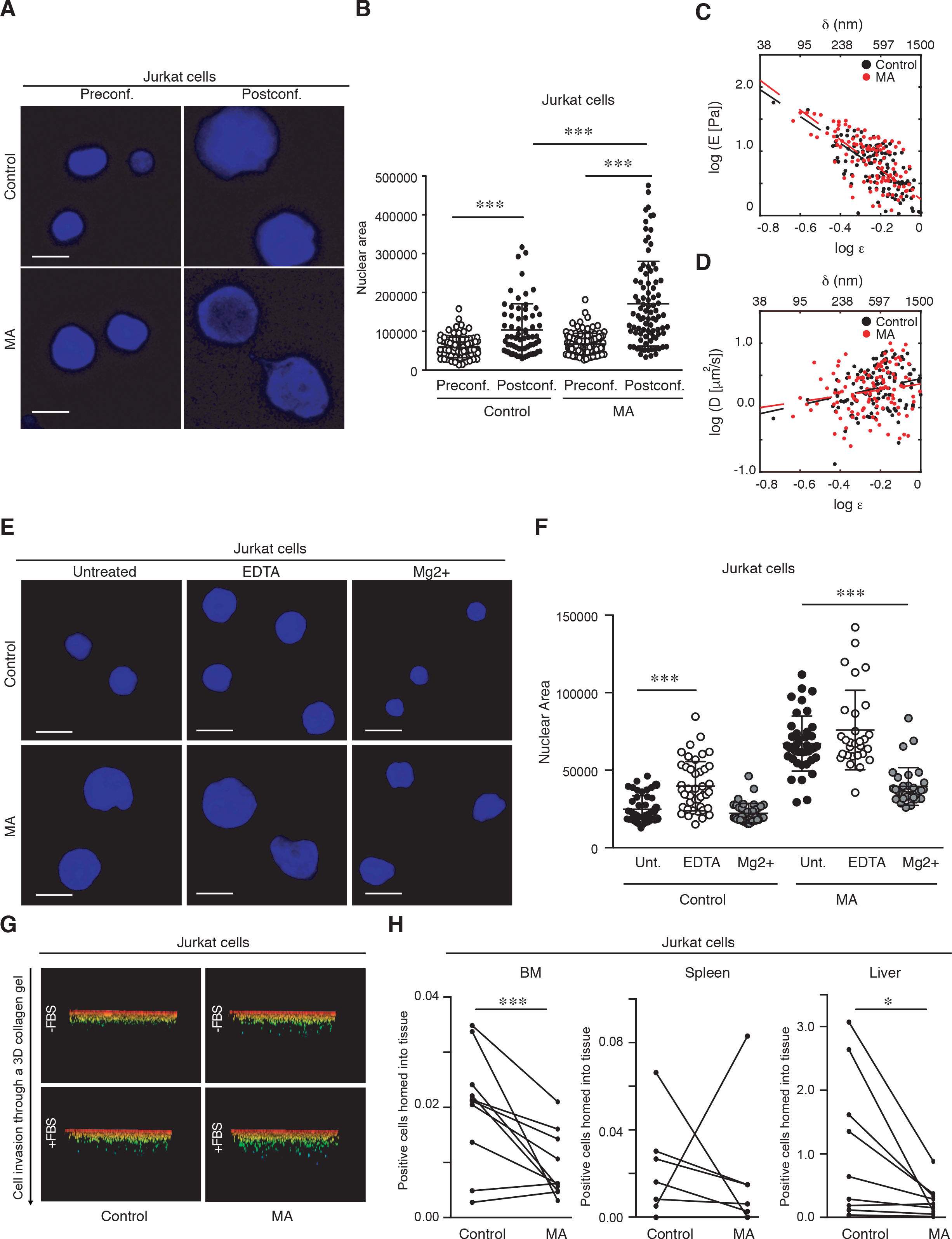
Migrated altered cells show altered mechanical and functional properties. **(A)** Isolated nuclei from control and MA Jurkat cells were stained with DAPI and seeded on poly-L-lysine-coated coverslips. Confocal sections of the nuclei were taken before (Preconf.) and after (Postconf.) confinement. Bar 10 μm. **(B)** Graph shows the nuclear area from cells in (A). Mean n = 66-89 isolated nuclei ± SD (3 replicates). **(C and D)** Variation of the stiffness (E) and the poroelastic diffusivity coefficient (D) in function of the nuclear indentation depth (shown in terms of strain, ε), both expressed as logarithmic values. Equivalent indentation depth in terms of distance, δ, is shown on top. Black and red dots represent values calculated from experimental data for control (n=124; indentation sites=13; nuclei=6) and MA Jurkat cells (n=115; indentation sites=12; nuclei=5), respectively. Linear fits are included as dashed lines. **(E)** Isolated nuclei from control and MA Jurkat cells were seeded on poly-lysine coated glasses. Then, osmotic stress conditions were induced by the addition of EDTA (swelling conditions) or MgCl_2_ (shrinking conditions). Nuclei were fixed, permeabilized and stained with DAPI for their analysis by confocal microscopy. Bar 10 μm. **(F)** Graph shows the nuclear area quantified from (C). Mean n = 29-46 isolated nuclei ± SD (2 replicates). **(G)** Control and MA Jurkat cells were seeded on the top of a collagen matrix and allowed to penetrate into the collagen in response to serum (FBS, fetal bovine serum) for 24h. Cells were fixed, stained with propidium iodide and serial confocal sections were captured. Images show the cell penetrability into the collagen. **(H)** Control (Cell Tracker Far Red+) and MA (CFSE+) Jurkat cells were mixed and injected into the tail vein of 10 NSG mice. After 24 h, labeled cells in spleen, liver and bone marrow were determined by flow cytometry. Graph shows the percentage of control and the MA cells analyzed in each animal. Mean n = 10.

As nuclear deformability is a critical stage for cell invasion [12], we assayed how permanent changes induced by constrained migration might affect this cellular function. First, we characterized *in vitro* that MA cells showed a higher chemotactic response compared to control cells (Figure S5A). Interestingly, MA cells showed a similar capacity to penetrate into a 3D collagen matrix as control cells (Figures 5G, and S5B). To determine the migratory capacity of MA cells and their ability to infiltrate into the bone marrow and spleen of immunodeficient mice [30], we injected an equal number of control and MA cells labeled with vital dyes into the tail vein of recipient mice and found a reduction of MA cells reaching these tissues related to harboring metastatic leukemia cells (Figures 5H, and S5C). These results indicate that constrained migration alters the mechanical behavior of the nucleus, which might have functional implications on the capacity of MA cells to infiltrate tissues.

### Actin homeostasis controls lamin B1 redistribution and the nuclear changes observed in migrating cells

Actin polymerization regulates the nuclear morphology and chromatin remodeling upon cellular constrictions [31, 32]. As we found that nuclear lamin B1 redistributed in MA cells, we used specific drugs to depolymerize or stabilize actin filaments. We observed that latrunculin (an inhibitor against actin polymerization) treatment was sufficient to rescue the localization of lamin B1 at the nuclear periphery in MA cells (Figures 6A, 6B, S6A, and S6B). Likely, MA cells treated with latrunculin showed a reduction in the nuclear area, whilst the pharmacological stabilization of polymerized actin with jasplakinolide did not promote any effect (Figure 6C). In addition to actin polymerization, several kinases trigger lamin B1 redistribution, such as Cdk1 (cyclin-dependent kinase 1) and protein kinase C (PKC) [33]. To determine whether these kinases might affect the phenotype observed in MA cells, we incubated these cells with specific inhibitors and observed that none of them rescued the normal lamin B1 redistribution (Figure S6C). Moreover, we did not see any reduction in the nuclear area of MA cells upon these treatments, and even the Cdk1 inhibitor (roscovitine) seemed to increase it (Figure S6D). Given that actomyosin controls the nuclear size and chromatin compaction [34, 35], we next investigated whether actin polymerization might also regulate the global the chromatin conformation in MA cells. We treated control and MA cells with DNAse I and found that MA cells were less sensitive to DNA digestion, suggesting a different global chromatin compaction. Furthermore, latrunculin treatment reverted this change in MA cells, which showed a DNA digestion profile more similar to control cells (Figures 6D, 6E, S6E and S6F). To further confirm this point, we performed *in situ* DNA digestion and observed that control cells were more sensitive to DNA digestion (visualized as a reduction in the nuclear area by DNAse activity) than MA cells (Figure S6G). Finally, several reports show that actin cytoskeleton is required during DNA damage response [36, 37]. We investigated the levels of γH2AX in MA cells treated with latrunculin and jasplakinolide, and found that γH2AX were reduced in MA cells treated with latrunculin, whereas jasplakinolide treatment exhibited a more similar phenotype to untreated MA cells (Figures 6F, and 6G). Taken together, our results indicate that actin polymerization regulates several of the permanent nuclear changes induced by persistent cell migration in MA cells.

**Figure 6:**
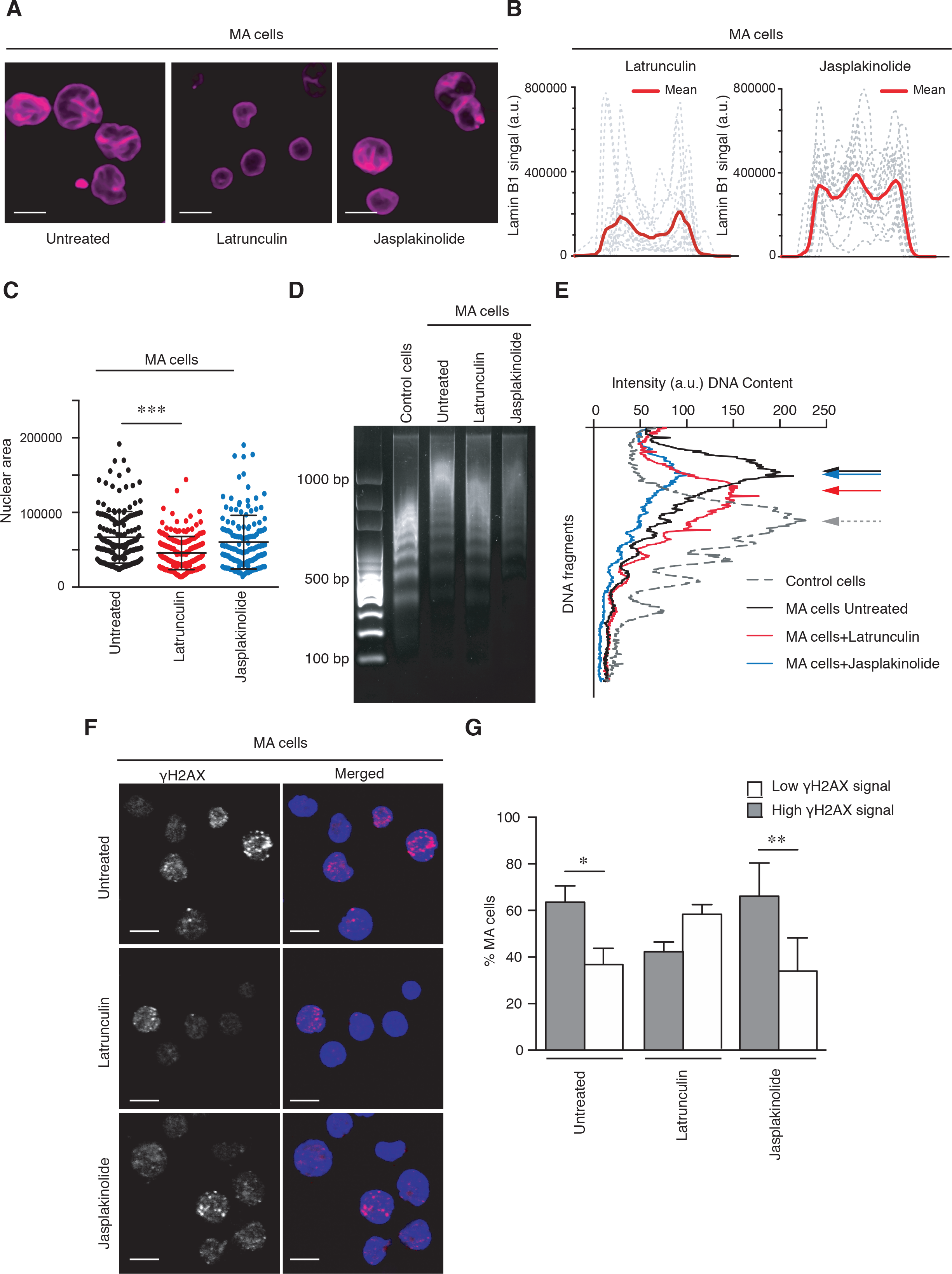
Actin polymerization balance regulates the nuclear changes described for MA cells. **(A)** MA Jurkat cells were cultured in the presence or absence of latrunculin B (1 μg/mL) and jasplakinolide (1 μg/mL) for 1h at 37°C. Then, cells were seeded on poly-L-lysine coated coverslips, fixed and analyzed by confocal microscopy. Bar 10 μm. **(B)** Line plots show the signal profile of lamin B1 from 15 representative nuclei. The red line indicates the mean intensity of the profiles analyzed. **(C)** Graph shows changes in the nuclear area of MA Jurkat cells under indicated treatments. Mean n= 145-185 nuclei ± SD (3 replicates). **(D)** Control and MA Jurkat cells were treated or not with latrunculin B or jasplakinolide for 1h. Then, cells were collected, and their DNA was digested with DNAse for 15 min and resolved in an agarose gel. **(E)** Graph shows the nucleosomal releasing profile from control (dashed line) and MA Jurkat cells as in (D). Arrows indicate the maxima DNA peaks in each cell population. **(F)** Control and MA Jurkat cells were treated or not with latrunculin B or jasplakinolide for 1h. Cells were fixed, permeabilized and stained with DAPI (blue) and γH2AX (red) for their analysis by confocal microscopy. Bar 10 μm. **(G)** Graph shows the percentage of cells with more than 2 visible foci for γH2AX from (F). Mean n= 53-82 nuclei ± SD (3 replicates).

## Discussion

In this work, we pointed out a connection between cell migration and permanent changes in the nucleus that lead to functional consequences in cell biology. The nucleus is a mechanosensory organelle that has to alter its mechanical and morphological characteristics for multiple cellular functions, including migration, development, and cancer progression [38, 39]. To undergo migration, cells have to sense their physiological environment to control their integrity and functions [40–42]. This is particularly important as extracellular migratory and mechanical signals regulate the adaptability of cells, their stemness, differentiation, genomic instability. Therefore, nuclear changes in response to cell migration might represent a major medical issue in many human diseases, including cancer infiltration [43–45].

Despite previous evidence showing that cell migration through confined spaces promotes nuclear ruptures, protein redistribution, and DNA damage nuclear alterations [46-48], their long-term consequences remain unclear. The impact of mechanobiology in stem cell reprogramming and cell rejuvenation has been previously described [49, 50], suggesting that cell confinement might influence memory effects and the fate transition of migratory cells. Our persistent cell migration approach shows that cells adapted to several rounds of cell migration by altering their nuclei in a permanent manner. This is particularly important as MA cells kept in culture showed a different phenotype and nuclear changes (including nuclear morphology and lamin B1 redistribution), which remain even without further cellular constriction. This transition highlights how cells might respond depending on the migratory stimulus and whether this is occasional or repeated. In addition, our data support that persistent migration controls altered transcription and phenotypical changes that agree with previous studies showing how biophysical stimuli control cell fate and differentiation [51, 52]. Mechanical stress between the cell and its environment is known to regulate cell cycle progression and genomic instability that leads hyperploidy [53, 54]. Although we did not see significant changes in cell proliferation, we cannot discard that under physiological conditions or external stimuli these changes might have potential implications in promoting cell state alterations and genomic instability and heterogeneity in cancer cells.

Due to the reported role of the nucleus as a mechanosensor [55], we analyzed the mechanical features of cells upon persistent and transient migration. To overcome the complexity of the mechanical analysis of isolated nuclei, we designed an experimental rationale based on nuclear swelling, nuclear compression, and indentation by optical tweezers. Previous works show nuclear swelling and compression as good experimental approaches to define the mechanical response of the nucleus [56-58]. In our experiments, the nucleus exhibits a two-regime response, depending on occasional or persistent cell migration. Considering the fact that migrating cells suffer different degree of mechanical stress on their nuclei; after a sole migration event, ORM cells might show some heterogeneity in their nuclear mechanical properties. On the other hand, MA cells could have developed a sort of homogeneous accommodation of the nuclear internal architecture due to the repeated application of a mechanical stress during cell migration, showing similar stiffness and poroelastic diffusivity. Permanent changes during the time of migratory cells may act in parallel or independently of other cellular functions. Interestingly, we described defective colonization of MA cells into different organs. This suggests that other molecular pathways could also be regulating the migration strategy used by MA cells, which might become less invasive than control leukemia cells. A plausible explanation might be a migratory exhaustion of MA cells, which would transit from a more effective movement into a more static condition. This versatility would be particularly important in several pathologies, including cancer, where invasive cells might switch into a heterogeneous population priming other cellular functions, such as cell cycle and resistance to apoptosis. Recent studies with normal and tumor cells have described the localization and upregulation of several markers of DNA damage in migratory cells under high nuclear deformation [59]. So far, nuclear lamins and deformability seemed to be involved in cell growth, DNA damage response, and cellular stress [16, 60]. From a clinical perspective, we pretended to confirm whether MA cells might show a therapeutic advantage in comparison to control leukemia cells. MA cells showed high basal levels of γH2AX but resisted apoptosis better than control cells. This chemoresistance to antimetabolites might be due to differences in cell proliferation or the metabolic activity of tumor cells [61]. Interestingly, we did not find changes in the proliferation of MA cells, which also show a downregulation of genes related to unfolded protein response and endoplasmic reticulum stress, fundamental in chemoresistance [62]. A plausible explanation is that under specific circumstances aberrant nuclear changes and high basal levels of γH2AX would be protective for MA cells against specific conventional therapies. This agrees with the idea that the steady state of γH2AX is critical for cell cycle regulation and the maintenance of genomic integrity [63, 64]. Our findings suggest that the potential adaptation of MA cells might have a dual effect on reducing their ability to reach other tissues, and on promoting other functions, such as cell survival and proliferation.

The cell skeleton, including actin polymerization and its associated proteins, is critical in many nuclear events, including chromatin compaction, and gene transcription. [65–67]. One exciting implication of our results is that actin depolymerization seemed to recapitulate the DNAse sensitivity of the chromatin in MA cells, in parallel to their lamin B1 redistribution at the nuclear periphery. Our observation also aligns with other results where actin homeostasis drugs alter the chromatin structure [68]. In fact, it has been reported that the actin cytoskeleton is critical for nuclear integrity and DNA repair in constricted migration [69]; and, in agreement with this, we observed that actin perturbation also reduced the levels of γH2AX in MA cells. Together, our findings highlight new knowledge about how cell migration underlies permanent nuclear changes that might lead to morphological, transcriptional, and functional changes in moving cells.

In conclusion, our findings highlight how migratory cells respond in structurally regulated ways to mechanical compression both in the short- and long-terms. We revealed a novel effect of persistent cell migration in morphological and transcriptional changes, which provides novel insights into alterations in the nucleus and functions of the cell. Furthermore, we have described how actin polymerization regulates the nuclear morphology and lamin B1 redistribution, supporting the relevance of molecular connections between the cell skeleton and nuclear changes induced by the cell environment interplay. Finally, we would like to propound the notion that cell migration underlies permanent nuclear changes that might lead to cell fate transition, genomic instability, and cellular heterogeneity with broad implications for development, aging, cancer, and mechanobiological processes.

## Materials and methods

### Cell lines and culture

The human Jurkat (CVCL_0367) and CCRF-CEM (RRID: CVCL_0207) cell lines were from ATCC, American Type Culture Collection. Both cell lines were monthly tested for mycoplasm contamination and maintained in culture in RPMI 1640 medium with L-glutamine and 125 μM HEPES (Sigma Aldrich, St. Louis, MO, USA) and 10% fetal bovine serum (Sigma-Aldrich) and maintained in 5% CO_2_ and 37°C.

One-round migration (ORM) cells were collected from the bottom chamber of Transwell inserts (Corning Costar, 6.5 mm diameter, 3 μm pore size) after 24h. For recovery assays, collected ORM cells were used to performed experiments (fresh condition); or kept in culture conditions for an additional 24h and then used to performed experiments (after 24h condition). Migratory-altered (MA) cells were generated as follows: Jurkat or CCRF-CEM cells were allowed to migrate in transwells for 24h; then, migrated cells were collected from the bottom chamber, counted and added again to the upper chamber of new inserts, under the same conditions. The same process was repeated 3 times. Finally, migrated cells from the bottom chamber were collected, expanded and kept in culture conditions as MA cells.

### Immunofluorescence

Cells were cultured in suspension, treated or not with specific inhibitors, for 1 hour at 37°C. Cells were seeded onto poly-L-lysine-coated glass slides for 30 min. Then, cells were fixed with 4% formaldehyde (15 min) and permeabilized with 0.5% Tx-100 in PBS (5 min). Samples were blocked in 10% fetal bovine serum with 0.1% Tx-100 in PBS, incubated with appropriated primary antibodies for 1h at RT, followed by several PBS washes and 1h at RT incubation with secondary antibodies. Samples were stained with DAPI 1 μg/mL for 10 min at RT, washed with PBS and H_2_O, and mounted. Images were acquired on an inverted DMi8 microscope (Leica) using an ACS-APO 63x NA 1.30 glycerol immersion objective. Quantification and analysis of images were determined using ImageJ.

### Electron microscopy

Cells were fixed for 1h in 3% glutaraldehyde in PBS and then washed twice with PBS. Samples were post-fixed in 1% osmium tetroxide and 0.8% potassium ferricyanide for 1h at 4°C and washed with PBS prior to dehydration with an increasing gradient of ethanol (30%, 50%, 70%, 80%, 90% and 100%) of 10 min per step. Samples were embedded in LX112 resin and were polymerized for 48h at 60°C. 60-80 nm sections were placed in copper grids of 75 mesh and stained with 5% uranyl acetate for 30 min and lead citrate for 4 min. Samples were viewed in a JEOL 1230 TEM and images were taken with a CMOS TVIPS 16 mp camera.

### Nuclear confinement

Cell nuclei were isolated by incubating cells in buffer A (10 μM HEPES, 10 μM KCl, 1.5 μM MgCl_2_, 0.34 M sucrose, 10% (v/v) glycerol, 1 μM DTT and Roche protease inhibitor cocktail) for 5 min at 4°C and centrifuged for 5 min at 4°C at 3,500 rpm. Nuclei were resuspended in TKMC buffer (50 μM Tris pH 7.5, 25 μM KCl, 3 μM MgCl_2_, 3 μM CaCl_2_), dyed with DAPI 1 μg/mL and sedimented onto poly-L-lysine coated plate for cell confinement analysis (4D cell). In this case, a glass slide with micropillars of 3 μm height was stuck on the silicone macropillar attached to the lid of the confiner. When the lid was closed, the pillars pushed the confining slides onto the culture substrate and confined the nuclei to 3 μm. Images of at least 20 nuclei were taken with a 63× objective before and after the confinement by an inverted confocal microscope.

### Flow cytometry

Isolated nuclei were resuspended in TKMC buffer and nuclear size was measured with a CytoFlex (Beckman Coulter). To analyze protein levels, cells were blocked with human IgG (1:1000; 30 min), incubated with specific primary antibodies of interest (1:1 to IgG), washed twice with PBS, followed by appropriated fluorochrome-conjugated secondary antibody for 30 min. Flow cytometry was performed with a FacSort. All data were analyzed using the Flow Jo software (Flow Jo LLC, Ashland, OR, USA).

### Optical tweezers

Isolated cell nuclei resuspended in TKM buffer were mixed with polystyrene beads with a mean particle size of 3.0 μm (Sigma-Aldrich) at final concentrations of 10^7^ nuclei/ml and 0.005% (w/v), respectively. The optical tweezers device (SensoCell, Impetux Optics S.L., Spain) is equipped with an ultra-stable single-frequency laser source (5 W, λ = 1064 nm) guided by acousto-optic deflectors and includes a direct force measurement platform capable of detecting the light scattered from the optical traps. The OT device is mounted on an inverted microscope (Eclipse Ti, Nikon, Japan) and a water immersion objective (Plan Apo VC 60XA/1.20 WI, Nikon) employed to focus the laser on the sample. A short-pass dichroic mirror transmits the bright-field illumination and reflects the IR trapping beam, and a short-pass filter was used to avoid IR laser radiation leaking. Sample bright-field imaging was captured by a CMOS camera (DCC1545M-GL, ThorLabs, USA) and the optical traps were operated with LightAce software (Impetux Optics S.L.). For each indentation experiment, a volume of 40 μL of the sample was placed in a custom-made glass chamber. This chamber was mounted on the microscope and measurements were performed at 25 °C. Only average-sized, round, symmetric nuclei with no alterations or major perturbations of their integrity were selected to perform the indentation routine. Indented nuclei got attached to the bottom surface of the glass chamber by unspecific interactions, and indenter beads were placed next to the nuclear envelope at axial positions and approximately 2 μm above the bottom surface of the chamber. Indenter beads had a nominal diameter of 3 μm, but the exact diameter of each bead used for indentation was measured by image analysis using the Fiji software. The indentation process consisted on pushing the nucleus laterally by generating an oscillation of the indenting bead. The fixed parameters of the oscillation are the shape (squared), frequency (0.5 Hz, which is enough to permit a complete relaxation between consecutive cycles), and offset (100%). The amplitude was variable, and the routine was set to sweep the 0.6-1.6 μm range with steps of 0.05 or 0.10 μm. Data were acquired during 45 s for each amplitude step. For each indentation experiment, the elasticity constant of the trap was calculated by using a particle scan routine included in the LightAce software. We wrote a specific Matlab (MathWorks, USA) script to analyze the data from the files obtained by the custom-made Labview indentation routine. This script is able to isolate each force-time curve for every indentation cycle and to calculate the stiffness from the indentation force values and the water diffusivity from the poroelastic relaxation times.

### Statistics

Statistical analysis and comparisons were made with GraphPad Prism6. The numerical data are presented as mean ± SD. Differences between means were tested by Student’s t test for two groups comparison. Where 3 or more groups were analyzed, one-way ANOVA was performed. P-values are indicated by asterisks ((*) P < 0.05; (**) P < 0.01; (***) P < 0.001).

## Declarations

### Ethics approval and consent to participate

AAll experiments involving animals were approved by the OEBA (Organ for Evaluating Animal Wellbeing) at CIB Margarita Salas and Madrid Regional Department of Environment, with reference PROEX 228.4/21; and carried out in strict accordance with the institution guidelines and the European and Spanish legislations for laboratory animal care.

### Consent for publication

All authors consent the manuscript for publication.

### Availability of data and materials

The accession numbers for the transcriptional microarray analyses is GSE214365. All relevant data are available from the corresponding author upon reasonable request.

### Authors’ contributions

A.dL.P., R.G.N., E.M., conducted experiments. H.Z.C., conducted experiments and contributed to data interpretation and discussion. V.T. contributed to data interpretation. H.L.M. conducted experiments and contributed to data interpretation. F.M., P.R.N., supervised the experiments, contributed to data interpretation and provided financing. J.R.M designed and supervised the experiments, contributed to data interpretation, wrote the paper, and provided financing.

## Acknowledgements

The authors thank the Microscopy Unit of Instituto de Investigación Biosanitaria Gregorio Marañón (IiSGM) for assistance with confocal analyses. The authors are also grateful to the EM and Animal Facilities of platforms of the CIB Margarita Salas for their assistance and support with the EM and *in vivo* experiments. The UCM-Genomic CAI Unit for their assistance with microarray experiments.

**Supplementary materials** include supporting material and methods, figures, tables and movies.

## References

1. te Boekhorst, V., Preziosi, P., and Friedl, P. (2016). Plasticity of cell migration in vivo and in silico. Annu. Rev. Cell Dev. Biol. 32, 491–526. doi: 10.1146/annurev-cellbio-111315-125201.

2. Ridle,y A.J., Schwartz, M.A., Burridge, K., Firtel, R.A., Ginsberg, M.H., Borisy, G., Parsons, J.T., and Horwitz, A.R. (2003). Cell migration: integrating signals from front to back. Science 302, 1704–1709. doi: 10.1126/science.1092053.

3. Paul, C.D., Shea, D.J., Mahoney, M.R., Chai, A., Laney, V., Hung, W.C., and Konstantopoulos, K. (2016). Interplay of the physical microenvironment, contact guidance and intracellular signaling in cell decision making. FASEB J. 30, 2161–2170. doi: 10.1096/fj.201500199R.

4. Vining, K.H., and Mooney, D.J. (2017). Mechanical forces direct stem cell behaviour in development and regeneration. Nat. Rev. Mol. Cell Bio. 18, 728–742. doi: 10.1038/nrm.2017.108.

5. Kirby, T.J., and Lammerding, J. (2018). Emerging views of the nucleus as a cellular mechanosensor. Nat. Cell Biol. 20, 373–381. doi: 10.1038/s41556-018-0038-y.

6. Burke, B., Stewart, C.L. (2014). Functional architecture of the cell’s nucleus in development, aging, and disease. Curr. Top Dev. Biol. 109, 1–52. doi: 10.1016/B978-0-12-397920-9.00006-8.

7. Gruenbaum, Y., and Foisner, R. (2015). Lamins: nuclear intermediate filament proteins with fundamental functions in nuclear mechanics and genome regulation. Annu. Rev. Biochem. 84, 131–164. doi: 10.1146/annurev-biochem-060614-034115.

8. Hübnet, M.R., and Spector, D.L. (2010). Chromatin dynamics. Annu. Rev. Biophys. 39, 471–489. doi: 10.1146/annurev.biophys.093008.131348.

9. Callan-Jones, A.. and Voituriez, R. (2016). Actin flows in cell migration: from locomotion and polarity to trajectories. Curr. Opin. Cell Biol. 38, 12–17. doi: 10.1016/j.ceb.2016.01.003.

10. Gupta, M., Sarangi, B.R., Deschamps, J., Nematbakhsh, Y., Callan-Jones, A., Margadant, F., Mège, R.M., Lim, C.T., Voituriez, R., and Ladoux, B. (2015). Adaptive rheology and ordering of cell cytoskeleton govern matrix rigidity sensing. Nat. Commun. 6, 7525. doi: 10.1038/ncomms8525.

11. Jain, N., Iyer, K.V., Kumar, A., and Shivashankar, G.V. (2013). Cell geometric constraints induce modular gene-expression patterns via redistribution of HDAC3 regulated by actomyosin contractility. Proc. Natl. Acad. Sci. USA 110, 11349–11354. doi: 10.1073/pnas.1300801110.

12. Denais, C.M., Gilbert, R.M., Isermann, P., McGregor, A.L., te Lindert, M., Weigelin, B., Davidson, P.M., Friedl, P., Wolf, K., and Lammerding, J. (2016). Nuclear envelope rupture and repair during cancer cell migration. Science 352, 353–358. doi: 10.1126/science.aad7297.

13. Raab, M., Gentili, M., de Belly, H., Thiam, H.R., Vargas, P., Jimenez, A.J., Lautenschlaeger, F., Voituriez, R., Lennon-Duménil, A.M., Manel, N., et al. (2016). ESCRT III repairs nuclear envelope ruptures during cell migration to limit DNA damage and cell death. Science 352, 359–362. doi: 10.1126/science.aad7611.

14. Irianto, J., Xia, Y., Pfeifer, C.R., Athirasala, A., Ji, J., Alvey, C., Tewari, M., Bennett, R.R., Harding, S.M., Liu, A.J., Greenberg, R.A., and Discher, D.E. (2017). DNA damage follows repair factor depletion and portends genome variation in cancer cells after pore migration. Curr. Biol. 27, 210–223. doi: 10.1016/j.cub.2016.11.049.

15. Pfeifer, C.R., Xia, Y, Zhu, K., Liu, D., Irianto, J., García, V.M.M., Millán, L.M.S., Niese, B., Harding, S., Deviri, D., et al. (2019). Constricted migration increases DNA damage and independently represses cell cycle. Mol. Biol. Cell 29, 1948–1962. doi: 10.1091/mbc.E18-02-0079.

16. Cho, S., Irianto, J., and Discher, D.E. (2017). Mechanosensing by the nucleus: From pathways to scaling relationships. J. Cell Biol. 216, 305–315. doi: 10.1083/jcb.201610042.

17. Ingber, D.E. (2013). Mechanobiology and diseases of mechanotransduction. Ann. Med. 35, 564–577. doi: 10.1080/07853890310016333.

18. Madrazo, E., González-Novo, R., Ortiz-Placín, C., García de Lacoba, M., Gonález-Murillo, Á., Ramírez, M., and Redondo-Muñoz, J. (2022). Fast H3K9 methylation promoted by CXCL12 contributes to nuclear changes and invasiveness of T-acute lymphoblastic leukemia cells. Oncogene 41, 1324–1336. doi: 10.1038/s41388-021-02168-8.

19. Harada, T., Swift, J., Irianto, J., Shin, J.W., Spinler, K.R., Athirasala, A., Diegmiller, R., Dingal, P.C., Ivanovska, I.L., and Discher, D.E. (2014). Nuclear lamin stiffness is a barrier to 3D migration, but softness can limit survival. J. Cell Biol. 204, 669–82. doi: 10.1083/jcb.201308029.

20. Uhler, C., and Shivashankar, G.V. (2017). Regulation of genome organization and gene expression by nuclear mechanotransduction. Nat. Rev. Mol. Cell Biol. 18, 717–727. doi: 10.1038/nrm.2017.101.

21. Le, H.Q., Ghatak, S., Yeung, C.Y., Tellkamp, F., Günschmann, C., Dieterich, C., Yeroslaviz, A., Habermann, B., Pombo, A., Niessen, C.M., et al. (2016). Mechanical regulation of transcription controls Polycomb-mediated gene silencing during lineage commitment. Nat. Cell Biol. 218, 864–875. doi: 10.1038/ncb3387.

22. Picariello, H.S., Kenchappa, R.S., Rai, V., Crish, J.F., Dovas, A., Pogoda, K., McMahon, M., Bell, E.S., Chandrasekharan, U., Luu, A., et al. (2019). Myosin IIA suppresses glioblastoma development in a mechanically sensitive manner. Proc. Natl. Acad. Sci. USA 116:15550–15559. doi: 10.1073/pnas.1902847116.

23. Cho, S., Vashisth, M., Abbas, A., Majkut, S., Vogel, K., Xia, Y., Ivanovska, I.L., Irianto, J., Tewari, M., Zhu, K., Tichy, E.D., et al. (2019). Mechanosensing by the lamina protects against nuclear rupture, DNA damage, and cell-cycle arrest. Dev. Cell 49, 920–935.e5. doi: 10.1016/j.devcel.2019.04.020.

24. Irianto, J., Xia, Y., Pfeifer, C.R., Greenberg, R.A., and Discher, D.E. (2017). As a nucleus enters a small pore, chromatin stretches and maintains integrity, even with DNA breaks. Biophys. J. 112, 446–449. doi: 10.1016/j.bpj.2016.09.047.

25. Akman, M., Belisario, D.C., Salaroglio, I.C., Kopecka, J., Donadelli, M., De Smaele, E., and Riganti C. (2021). Hypoxia, endoplasmic reticulum stress and chemoresistance: dangerous liaisons. J. Exp. Clin. Cancer Res. 40, 28. doi: 10.1186/s13046-020-01824-3.

26. Toksvang, L.N., Lee, S.H.R., Yang, J.J., and Schmiegelow, K. (2022). Maintenance therapy for acute lymphoblastic leukemia: basic science and clinical translations. Leukemia 36, 1749–1758. doi: 10.1038/s41375-022-01591-4.

27. Stephens, A.D., Liu, P.Z., Banigan, E.J., Almassalha, L.M., Backman, V., Adam, S.A., Goldman, R.D., and Marko, J.F. (2018). Chromatin histone modifications and rigidity affect nuclear morphology independent of lamins. Mol. Biol. Cell 29, 220–233. doi: 10.1091/mbc.E17-06-0410.

28. Le Berre, M., Aubertin, J., and Piel, M. (2012). Fine control of nuclear confinement identifies a threshold deformation leading to lamina rupture and induction of specific genes. Integr. Biol. 4, 1406–1414. doi: 10.1039/c2ib20056b.

29. Finan, J.D., and Guilak, F. (2010). The effects of osmotic stress on the structure and function of the cell nucleus. J. Cell Biochem. 109, 460–467. doi: 10.1002/jcb.22437.

30. Valent, P., Sadovnik, I., Eisenwort, G., Herrmann, H., Bauer, K., Mueller, N., Sperr, W.R., Wicklein, D., and Schumacher U. (2020). Redistribution, homing and organ-invasion of neoplastic stem cells in myeloid neoplasms. Semin. Cancer Biol. 60, 191–201. doi: 10.1016/j.semcancer.2019.07.025.

31. Croft, D.R., Coleman, M.L., Li, S., Robertson, D., Sullivan, T., Stewart, C.L., and Olson, M.F. (2005). Actin myosin-based contraction is responsible for apoptotic nuclear disintegration. J. Cell Biol. 168, 245–255. doi: 10.1083/jcb.200409049.

32. Hatch, M.E., and Hetzer, M.W. (2016). Nuclear envelope rupture is induced by actin-based nucleus confinement. J. Cell Biol. 215, 27–36. doi: 10.1083/jcb.201603053.

33. Mall, M., Walter, T., Gorjánácz, M., Davidson, I.F., Nga Ly-Hartig, T.B., Ellenberg, J., and Mattaj, I.W. (2012). Mitotic lamin disassembly is triggered by lipid-mediated signaling. J. Cell Biol. 198, 981–990. doi: 10.1083/jcb.201205103.

34. Takaki, T., Montagner, M., Serres, M.P., Le Berre, M., Russell, M., Collinson, L., Szuhai, K., Howell, M., Boulton, S.J., Sahai, E., and Petronczki, M. (2017). Actomyosin drives cancer cell nuclear dysmorphia and threatens genome stability. Nat Comms. 2017;8:16013. doi: 10.1038/ncomms16013.

35. Buxboim, A., Swift, J., Irianto, J., Spinler, K.R., Dingal, P.C., Athirasala, A., Kao, Y.R., Cho, S., Harada, T., Shin, J.W., and Discher, D.E. (2014). Matrix elasticity regulates lamin-A,C phosphorylation and turnover with feedback to actomyosin. Curr. Biol. 24, 1909–17. doi: 10.1016/j.cub.2014.07.001.

36. Hurst, V., Challa, K., Shimada, K., and Gasser, S.M. (2021). Cytoskeleton integrity influences XRCC1 and PCNA dynamics at DNA damage. Mol. Biol. Cell 32, br6. doi: 10.1091/mbc.E20-10-0680.

37. Wang, L., Wang, M., Wang, S., Qi, T., Guo, L., Li, J., Qi, W., Ampah, K.K., Ba, X., and Zeng, X. (2013). Actin polymerization negatively regulates p53 function by impairing its nuclear import in response to DNA damage. PLoS One 8, e60179. doi: 10.1371/journal.pone.0060179.

38. Dahl, K.N., Ribeiro, A.J.S., and Lammerding, J. (2008). Nuclear shape, mechanics, and mechanotransduction. Circ. Res. 102, 1307–1318. doi: 10.1161/CIRCRESAHA.108.173989.

39. Zwerger, M., Ho, C.Y., and Lammerding, J. (2011). Nuclear mechanics in disease. Annu. Rev. Biomed. Eng. 13, 397–428. doi: 10.1146/annurev-bioeng-071910-124736.

40. Charras, G., and Sahai, E. (2014). Physical influences of the extracellular environment on cell migration. Nat. Rev. Mol. Cell Biol. 15, 813–824. doi: 10.1038/nrm3897.

41. Yamada, K.M., Collins, J.W., Cruz Walma, D.A., Doyle, A.D., Morales, S.G., Lu, J., Matsumoto, K., Nazari, S.S., Sekiguchi, R., Shinsato, Y., and Wang, S. (2019). Extracellular matrix dynamics in cell migration, invasion and tissue morphogenesis. Int. J. Exp. Path. 100, 144–152. doi: 10.1111/iep.12329.

42. Liu, Y.J., Le Berre, M., Lautenschlaeger, F., Maiuri, P., Callan-Jones, A., Heuzé, M., Takaki, T., Voituriez, R., and Piel, M. (2015). Confinement and low adhesion induce fast amoeboid migration of slow mesenchymal cells. Cell 160, 659–672. doi: 10.1016/j.cell.2015.01.007.

43. Yang, C., Del Rio, F.W., Ma, H., Killaars, A.R., Basta, L.P., Kyburz, K.A., and Anseth, K.S. (2016). Spatially patterned matrix elasticity directs stem cell fate. Proc. Natl. Acad. Sci. USA 113, E4439–E4445. doi: 10.1073/pnas.1609731113.

44. Heo, S.J., Han, W.M., Szczesny, S.E., Cosgrove, B.D., Elliott, D.M., Lee, D.A., Duncan, R.L., and Mauck, R.L. (2016). Mechanically induced chromatin condensation requires cellular contractility in mesenchymal stem cells. Biophys. J. 111, 864–874. doi: 10.1016/j.bpj.2016.07.006.

45. Stefanello, S.T., Luchtefeld, I., Liashkovich, I., Pethö, Z., Azzam, I., Bulk, E., Rosso, G., Döhlinger, L., Hesse, B., Oeckinghaus, A., Shahin, V. (2021). Impact of the Nuclear Envelope on Malignant Transformation, Motility, and Survival of Lung Cancer Cells. Adv Sci (Weinh). 8, e2102757. doi: 10.1002/advs.202102757.

46. Xia, Y., Pfeifer, C.R., Zhu, K., Irianto, J., Liu, D., Pannell, K., Chen, E.J., Dooling, L.J., Tobin, M.P., Wang, M,, et al. (2019). Rescue of DNA damage after constricted migration reveals a mechano-regulated threshold for cell cycle. J. Cell Biol. 218, 2545–2563. doi: 10.1083/jcb.201811100.

47. Irianto, J., Pfeifer, C.R., Bennett, R.R., Xia, Y., Ivanovska, I.L., Liu, A.J., Greenberg, R.A., and Discher, D.E. (2017). Nuclear constriction segregates mobile nuclear proteins away from chromatin. Mol. Biol. Cell 27, 4011–4020. doi: 10.1091/mbc.E16-06-0428.

48. Shah, P., Hobson, C.M., Cheng, S., Colville, M.J., Paszek, M.J., Superfine, R., and Lammerding, J. (2021). Nuclear deformation causes DNA damage by increasing replication stress. Curr. Biol. 31, 1–13. doi: 10.1016/j.cub.2020.11.037.

49. Engler, A.J., Sen, S., Sweeney, H.L., and Discher, D.E. (2006). Matrix elasticity directs stem cell lineage specification. Cell 126, 677–689. doi: 10.1016/j.cell.2006.06.044.

50. Killaars, A.R., Grim, J.C., Walker, C.J., Hushka, E.A., Brown, T.E., and Anseth, K.S. (2018). Extended exposure to stiff microenvironments leads to persistent chromatin remodeling in human mesenchymal stem cells. Adv. Sci. (Weinh) 6, 1801483. doi: 10.1002/advs.201801483.

51. Downing, T.L., Soto, J., Morez, C., Houssin, T., Fritz, A., Yuan, F., Chu, J., Patel, S., Schaffer, D.V., and Li, S. (2013). Biophysical regulation of epigenetic state and cell reprogramming. Nat. Mater. 12, 1154–1162. doi: 10.1038/nmat3777.

52. Werner, M., Blanquer, S.B., Haimi, S.P., Korus, G., Dunlop, J.W., Duda, G.N., Grijpma, D.W., Petersen, A. (2016). Surface Curvature Differentially Regulates Stem Cell Migration and Differentiation via Altered Attachment Morphology and Nuclear Deformation. Adv Sci (Weinh). 4, 1600347. doi: 10.1002/advs.201600347.

53. Uroz, M., Wistorf, S., Serra-Picamal, X., Conte, V., Sales-Pardo, M., Roca-Cusachs, P., and Trepat, X. (2018). Regulation of cell cycle progression by cell-cell and cell-matrix forces. Nat. Cell Biol. 20, 461–654. doi: 10.1038/s41556-018-0107-2.

54. Gemble, S., Wardenaar, R., Keuper, K., Srivastava, N., Nano, M., Macé, A.S., Tijhuis, A.E., Bernhard, S.V., Spierings, D.C.J., Simon, A., et al. Genetic instability from a single S phase after whole-genome duplication. Nature. 2018, 604:146–151. doi: 10.1038/s41586-022-04578-4.

55. Guilluy, C., Osborne, L.D., Van Landeghem, L., Sharek, L., Superfine, R., Garcia-Mata, R., and Burridge, K. (2014). Isolated nuclei adapt to force and reveal a mechanotransduction pathway in the nucleus. Nat. Cell Biol. 16, 376–381. doi: 10.1038/ncb2927.

56. Furusawa, T., Rochman, M., Taher, L., Dimitriadis, E.K., Nagashima, K., Anderson, S., and Bustin, M. (2015). Chromatin decompaction by the nucleosomal binding protein HMGN5 impairs nuclear sturdiness. Nat. Commun. 6, 6138. doi: 10.1038/ncomms7138.

57. Zhang, X., Cook, P.C., Zindy, E., Williams, C.J., Jowitt, T.A., Streuli, C.H., MacDonald, A.S., and Redondo-Muñoz, J. (2016). et al. Integrin α4β1 controls G9a activity that regulates epigenetic changes and nuclear properties required for lymphocyte migration. Nucleic Acids Res. 44, 3031–44. doi: 10.1093/nar/gkv1348.

58. Wang, P., Dreger, M., Madrazo, E., Williams, C.J., Samaniego, R., Hodson, N.W., Monroy, F., Baena, E., Sánchez-Mateos, P., Hurlstone, A., et al. (2020). WDR5 modulates cell motility and morphology and controls nuclear changes induced by a 3D environment. Proc Natl Acad Sci USA. 2020;115:8581–8586. doi: 10.1073/pnas.1719405115.

59. Kumar, A., Mazzanti, M., Mistrik, M., Kosar, M., Beznoussenko, G.V., Mironov, A.A., Garrè, M., Parazzoli, D., Shivashankar, G.V., Scita, G., et al. (2014). ATR mediates a checkpoint at the nuclear envelope in response to mechanical stress. Cell 158, 633–646. doi: 10.1016/j.cell.2014.05.046.

60. Aureille, J., Buffiere-Ribot, V., Harvey, B.E., Boyault, C., Pernet, L., Andersen, T., Bacola, G., Balland, M., Fraboulet, S., Van Landeghem, L., and Guilluy, C. (2019). Nuclear envelope deformation controls cell cycle progression in response to mechanical force. Embo Rep. 20, e48084. doi: 10.15252/embr.201948084.

61. Karran, P., and Attard, N. (2008). Thiopurines in current medical practice: molecular mechanisms and contributions to therapy-related cancer. Nat. Rev. Cancer. 8, 24–36. doi: 10.1038/nrc2292.

62. Oakes, S.A. (2020). Endoplasmic reticulum stress signaling in cancer cells. Am. J. Pathol. 190, 934–946. doi: 10.1016/j.ajpath.2020.01.010.

63. Meador, J.A., Zhao, M., Su, Y., Narayan, G., Geard, C.R., and Balajee, A.S. (2008). Histone H2AX is a critical factor for cellular protection against DNA alkylating agents. Oncogene 27, 5662–5671. doi: 10.1038/onc.2008.187.

64. Celeste, A., Petersen, S., Romanienko, P.J., Fernandez-Capetillo, O., Chen, H.T., Sedelnikova, O.A., Reina-San-Martin, B., Coppola, V., Meffre, E., Difilippantonio, M.J., et al. (2002). Genomic instability in mice lacking histone H2AX. Science 296, 922–927. doi: 10.1126/science.1069398.

65. Tusamda Wakhloo, N., Anders, S., Badique, F., Eichhorn, M., Brigaud, I., Petithory, T., Vassaux, M., Milan, J.L., Freund, J.N., Rühe, J., et al., (2020). Actomyosin, vimentin and LINC complex pull on osteosarcoma nuclei to deform on micropillar topography. Biomaterials 234, 119746. doi: 10.1016/j.biomaterials.2019.119746.E

66. Schrank, B.R., Aparicio, T., Li, Y., Chang, W., Chait, B.T., Gundersen, G.G., Gottesman, M.E., and Gautier, J. (2018). Nuclear Arp2/3 drives DNA break clustering for homology-directed repair. Nature 559, 61–66. doi: 10.1038/s41586-018-0237-5

67. Mistriotis, P., Wisniewski, E.O., Bera, K., Keys, J., Li, Y., Tuntithavornwat, S., Law, R.A., Perez-Gonzalez, N.A., Erdogmus, E., Zhang, Y., Zhao, R., et al. (2019). Confinement hinders motility by inducing RhoA-mediated nuclear influx, volume expansion and blebbing. J. Cell Biol. 218, 4093–4111. doi: 10.1083/jcb.201902057.

68. Makhija, E., Jokhun, D.S., and Shivashankar, G.V. (2016). Nuclear deformability and telomere dynamics are regulated by cell geometric constraints. Proc. Natl. Acad. Sci. USA 113, E32–E40. doi: 10.1073/pnas.1513189113.

69. Patteson, A.E., Vahabikashi, A., Pogoda, K., Adam, S.A., Mandal, K., Kittisopikul, M., Sivagurunathan, S., Goldman, A., Goldman, R.D., and Janmey, P.A. (2019). Vimentin protects cells against nuclear rupture and DNA damage during migration. J. Cell Biol. 218, 4079–4092. doi: 10.1083/jcb.201902046.

