## Supplemental Info for "Persistent confined migration confers permanent nuclear and functional changes in migrating cells"

**Supplementary Materials and Methods.**

**Supplementary Figures S1-S6.**

**Supplementary Tables S1 and S2.**

**Supplementary Movies S1-S4.**

#### SUPPLEMENTARY MATERIALS AND METHODS

| Antibodies & reagents | Conc use | Host | Company | #Cat |
|---|---|---|---|---|
| DAPI | 1:1000 | - | Sigma-Aldrich | 62248 |
| Anti-lamin B1 | 1:100 | Mouse | Santa Cruz | 365962 |
| Anti-pH2AX | 1:100 | Mouse | Invitrogen | 14-9865-82 |
| Anti-PKC-α | 1:1000 | Rabbit | Invitrogen | PA5-85652 |
| Anti-PKC-β | 1:1000 | Rabbit | Thermo Fisher | PA5-13740 |
| Anti-WDR5 | 1:1000 | Rabbit | Bethyl | A302-430-A-M |
| Anti-EZH2 | 1:1000 | Rabbit | Cell Signaling | 5246S |
| Anti-SYK | 1:1000 | Rabbit | Cell Signalling | D3Z1E |
| Anti-β-actin | 1:100 | Rabbit | Sigma-Aldrich | SAB5600204 |
| CF647- anti-mouse IgG (H+L) | 1:100 | Donkey | Biotium | 20042 |
| CF594- anti-rabbit IgG (H+L) | 1:100 | Donkey | Biotium | 20152 |
| Anti-rabbit HRP- conjugated | 1:5000 | Goat | BioRad | 170-6515 |
| Anti-mouse HRP- conjugated | 1:5000 | Goat | BioRad | 170-6516 |
| CellTrace™ CFSE | 1 µM | - | Thermo Scientific | C34554 |
| CellTrace™ Far Red | 1 µM | - | Thermo Scientific | C34564A |
| Poly-L-Lysine | - | - | Sigma-Aldrich | 25988-63-0 |
| DACO | - | - | Sigma-Aldrich | 10981 |
| Propidium iodide | 50 µg/ml | | Sigma-Aldrich | P4864 |
| MTT | 1:10 | - | Sigma-Aldrich | CT01-5 |
| Methotrexate | 1 µM | - | Sigma-Aldrich | M9929 |
| Proteinase K | 1 µg/mL | - | Sigma-Aldrich | P1308 |
| RNAse A | 100 µg/ml | - | Sigma-Aldrich | R4875 |

| | | | | |
|---|---|---|---|---|
| Roscovitine | 15 µM | - | Sigma-Aldrich | R7772 |
| Staurosporine | 50 nM | - | Sigma-Aldrich | 19-123-M |
| Enzastaurin | 2 nM | - | Sigma-Aldrich | SML0762 |
| DNAse I | 1U or 50U | - | Thermo Fisher | EN0521 |
| 35mm glass-bottomed plates | - | - | Ibidi | 80136 |
| Phenol:chloroform:isoamyl alcohol | - | - | Panreac | A0889.0250 |
| Latrunculin B | 1 µg/mL | - | Enzo | BMC-T110_0001 |
| Jasplakinolide | 1 µg/mL | - | Enzo | ALX-350-275-C050 |

**Immunoblotting.** Cells were washed with cold PBS, lysed in sample buffer and then sonicated (15 s at 70% amp) using a Microson XL2000 (Misonix). Protein samples were boiled and resolved by 7.5-15% SDS-PAGE and transferred to nitrocellulose membranes (GE Healthcare Life Science). After electrophoresis, membranes were with 5% low fat milk in TBS-Tween (0.5%) for 1h at RT and incubated overnight with primary antibodies (1:1000) at 4ºC with rotation. After washing with TBS-Tween 1%, membranes were incubated with HRP-labelled secondary antibodies (1:5000) for 1 hour at room temperature. Protein signal was developed using the enhanced chemiluminiscent detection method (Amersham) and analyzed in a ChemiDoc (Bio-Rad). For the stripping process, membranes were washed 3 times with stripping buffer (1.5% Glycine, 1% SDS and 1% NP-40 pH=2.2) for 1h each wash, before blocking and incubating with the proper primary antibodies as described above. Quantification and analysis of images was performed using ImageJ.

**Real-Time PCR (qPCR).** Total RNA was extracted using TRI reagent (Sigma) and genomic DNA was digested with 1U DNAse I, 2.5 mM MgCl2 (Promega) at 37ºC for 30 min. 5mM

EDTA and 10 min at 65ºC was used to stop digestion. 1 μg of purified RNA was retrotranscribed using First Strand cDNA synthesis kit (Thermofisher) and cDNA concentrations were quantified using a NanoDrop ND-1000 Spectrophotometer (Fisher Scientific). Oligonucleotides for selected genes were designed according to the PrimerQuest Tool (IDT): *JAK2:* 5′-GCAACAGGAACAAGATGTGAAC-3′ and 5′-TTCCCTCCATTTCTGTCATCG-3′; *EZH2*: 5’-TTTCCAACACAAGTCATCCC-3’ and 5’-AACCCACATTCTTATCCCC-3; and *qTBP:* 5′-CGGCTGTTTAACTTCGCTTC-3′ and 5′-CACACGCCAAGAAACAGTGA-3′. Quantitative real-time PCR (qRT-PCR) was performed on a Roche LightCycler 96 following the manufacturer's instructions. Assays were made in triplicates and results normalized according to the expression levels of TBP. Melt curve analysis was performed at the end of PCR to confirm the presence of a single, specific product. The results were expressed using the ΔΔCt method for quantification.

**RNA microarray**. The mRNAs from cells were isolated using the NucleoSpin RNA kit (Macherey-Nagel), according to the manufacturer’s instructions. The RNA purity and concentration were determined by Nanodrop measurement, and 1 μg of RNA was used for microarray analysis by Human Gene Clariom S Assay (Thermo Fisher Scientific). Data were processed, normalized and log2 transformed by the UCM-Genomic CAI Unit. Analysis was performed using Transcriptome Analysis Console and Database for Annotation, Visualization and Integrated Discovery (DAVID) v6.8. Microarray data set used in this study is deposited at GEO (accession number #GSE214365).

**Osmotic stress**. Isolated nuclei were resuspended in TKMC buffer and sedimented onto poly-L-lysine coated plates. Nuclei were incubated or not with 5 μM EDTA (swelling condition) or $MgCl_2$ (shrinking condition) for 10 min. Then, nuclei were fixed, permeabilized, and stained for their visualization on the microscope. Quantification and analysis of images were determined using ImageJ software.

**Transwell invasion**. Chemotaxis experiments were carried out using Transwell inserts (Corning Costar, 6.5 mm diameter, 3 μm pore size). 100 μL of serum-free RPMI containing $2x10^5$ ALL cells under specific conditions were added to the upper chamber of the insert, and 600 μL of RPMI supplemented with serum to the bottom chamber. After 24h, migrated cells were collected from the bottom chamber and counted to calculate migration index.

**Cell penetration assay**. A 100 μl collagen matrix was reconstituted at 1.7 mg/mL in RPMI, neutralized with 7.5% $NaHCO_3$ and 25 mM HEPES inside Transwell inserts (0.4 μm, Costar). After 1 h at 37ºC, 100 μL of serum-free RPMI containing $3x10^5$ cells was added on the top of it. RPMI medium with 10% of FBS was added to the bottom chamber of the Transwell as chemoatractant. After 24 h, invading cells were fixed with 4% PFA for 1 h, permeabilized with 0.5% Triton-X-100 in PBS for 30 min and stained with propidium iodide. Invading cells were imaged with a sCMOS Orca-Flash 4.0LT camera (Hamamatsu) coupled 5 to an inverted DMi8 microscope (Leica), capturing serial z- stacks every 10 μm with a 10× objective (dry ACS APO 10x/NA 0.3).

**In-vivo invasiveness**. NOD-SCID-Il2rg-/- (NSG) mice (Mus musculus), were bred and maintained at the Servicio del Animalario del Centro de Investigaciones Biológicas Margarita Salas (CIB-CSIC) with number 28079-21A. All mice were used following guidelines issued by the European and Spanish legislations for laboratory animal care. $5x10^6$ cells were labeled with Far Red Cell Tracker (1 μM, control cells) and CFSE (1 μM, MA cells) for 30 min. Then, cell populations were mixed and intravenously (IV) administered to 15 weeks-old non-conditioned NSG mice. Sacrifice was performed 24 hours after injection. Bone marrow from femurs, the spleen and the liver were extracted and processed through mechanical disaggregation. The resulting tissues and the peripheral blood were processed with RBC lysis buffer (Thermo Scientific), filtered with a 100 μm strainer (ClearLine), and analyzed by flow citometry (FACSCantoTM II, Becton Dickinson).

**MTT proliferation assay.** 1x$10^5$ cells were seeded into microplates in 100 µl culture medium for 24 or 48 h. Then, 10 µl of the MTT labeling reagent was added, and the microplate was kept for 4 h at 37°C in a humidified atmosphere. MTT was solubilized using 100 µL of isopropyl alcohol and 20 µL of PBS-3% SDS. Absorbance was measured at 560 nm (Varioskan, Thermo Fisher).

**BrdU cell proliferation assay**. 2x$10^4$ cells per condition were collected and then processed with the BrdU cell Proliferation Assay Kit (Cell Signaling) following the manufacturer's instructions. Briefly, after the BrdU addition the cells were incubated for either 4h or 18h at 37ºC to allow them to replicate. Then the cells were fixed and stained, first with a primary antibody for 1h at RT and then with an HRP-conjugated secondary antibody for 30 min at RT. Finally, HRP substrate TMB was added to develop color, and the absorbance was measured at 450 nm (Varioskan, Thermo Fisher).

**Cell cycle.** 3x$10^5$ cells were fixed in ice-cold 70% (v/v) ethanol overnight. Then, cells were centrifuged and the pellet was incubated in PBS with RNAse (100 µg/ml), 0.1% Triton X-100 and propidium iodide (50 µg/ml) for 30 min at RT. Cells were washed, resuspended in 600 µl of PBS-EDTA 2 mM and analyzed by flow cytometry. Data analysis was performed using the FlowJo software and the cell cycle distribution determined by G1, S, and G2/M cell populations.

**Cell viability.** Cells were treated with or without methotrexate (1 µM) for 24h, collected, washed in PBS and stained using the Annexin V-FITC Apoptosis Detection kit (Immunostep) according to manufacturer's instructions. The number of living cells was determined by Flow Cytometry, and data analysis was performed using the software FlowJo.

**DNAse I-sensitivity assay.** Cells were resuspended in Lysis buffer (Tris-HCl 10 µM [pH 7.5], sucrose 300 mM, NaCl 15 µM, $MgCl_2$ 5 µM, NP-40 0.5%, DTT 0.5 µM) with protease inhibitors. Then, lysates were digested with 1 U of DNAse I diluted in DNAse buffer (Tris-

HCl 10 μM [pH 7.5], $MgCl_2$ 2.5 μM, $CaCl_2$ 0.5 μM) for 8 or 15 min at 25°C. Reactions were stopped by adding STOP buffer (Tris-HCl 10 μM [pH 8], EDTA 5 μM, NaCl 200 μM, SDS 0.2%), and samples were subsequently incubated with RNAseA for 30 min at 37°C and proteinase K 60 min at 52°C. Digested DNA was washed with phenol/chloroform and precipitated with 2.5 volumes of ethanol and 3 M sodium acetate at −80°C overnight. DNA pellet was dissolved in water and resolved in 1.5% agarose gel.

**In situ DNAse assay**. Cells were seeded onto poly-L-lysine-coated glasses for 15 min at 37ºC and treated with CSK buffer (PIPES/KOH 10mM, NaCl 100mM, sucrose 300mM, $MgCl_2$ 1mM) for additional 5 min at RT. Then, cells were incubated with DNAse I (50U/mL) for 20 min. Remaining nuclei were clarified and stained with CSK buffer plus 125 mM $(NH_4)_2SO_4$ and Hoechst 33342 for 5 min at RT. Then, nuclei were fixed with methanol at −20°C for 5 min, washed with CSK buffer and mounted. Images were acquired on an inverted DMi8 microscope (Leica) using an ACS-APO 63x NA 1.30 glycerol immersion objective. Quantification and analysis of images were determined using ImageJ.

### SUPPLEMENTARY FIGURES

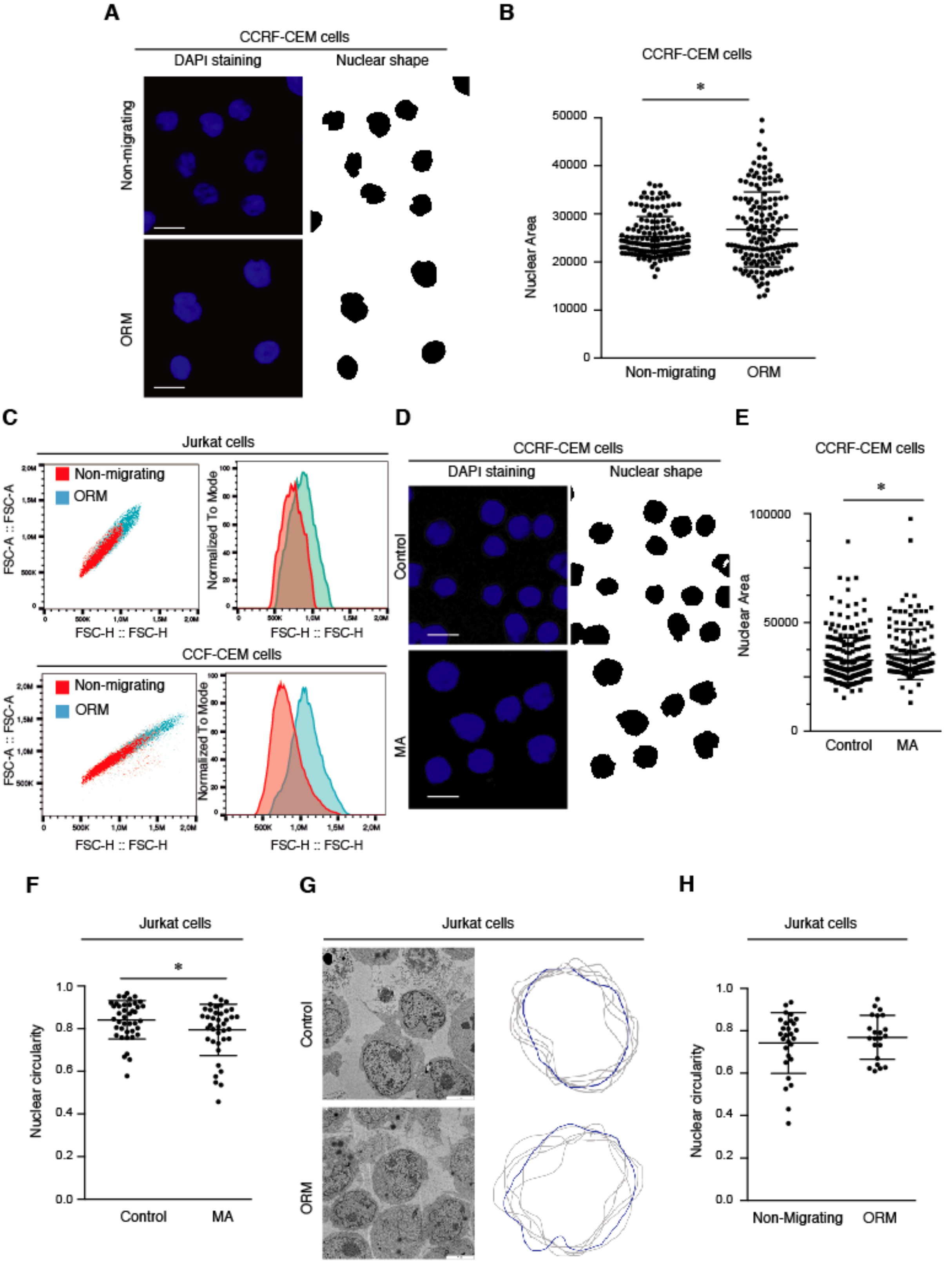


**Figure S1**

**Fig. S1.** **(A)** CCRF-CEM cells were allowed to migrate across 3 μm Transwell inserts for 24h. Non-migrating and one-round migrated (ORM) cells were collected from the upper and bottom chambers, respectively, sedimented on poly-L-lysine coated coverslips, fixed and stained with DAPI. Bar 10 μm. **(B)** Graph shows changes in the nuclear area of CCRF-CEM cells upon one round of migration through constrictions. Mean n= 157 cells ± SD (3 replicates). **(C)** Isolated nuclei from non-migrating and ORM cells were stained with DAPI, and their morphology and size were determined by flow cytometry. Graphs show the size of nuclei according to the FSC parameter. **(D)** Control and MA CCRF-CEM cells were sedimented on poly-L-lysine coated coverslips, fixed and stained with DAPI. Right panels indicate in black the area of the nuclei. Bar 10 μm. **(E)** Graph shows changes in the nuclear area of control and MA cells. Mean n= 161-258 cells ± SD (3 replicates). **(F)** Graph shows changes in the nuclear circularity of control and MA Jurkat cells quantified from images obtained in thin section electron microscopy. Mean n=39-41 cells ± SD. **(G)** Non-migrating and ORM Jurkat cells were collected and processed for thin section electron microscopy to visualize the nuclear morphology. Plots show changes in the nuclear circularity of non-migrating and ORM Jurkat cells (n=6 representative cells). **(H)** Graph shows changes in the nuclear circularity of non-migrating and ORM Jurkat cells quantified from images obtained in thin section electron microscopy. Mean n=22-28 cells ± SD.

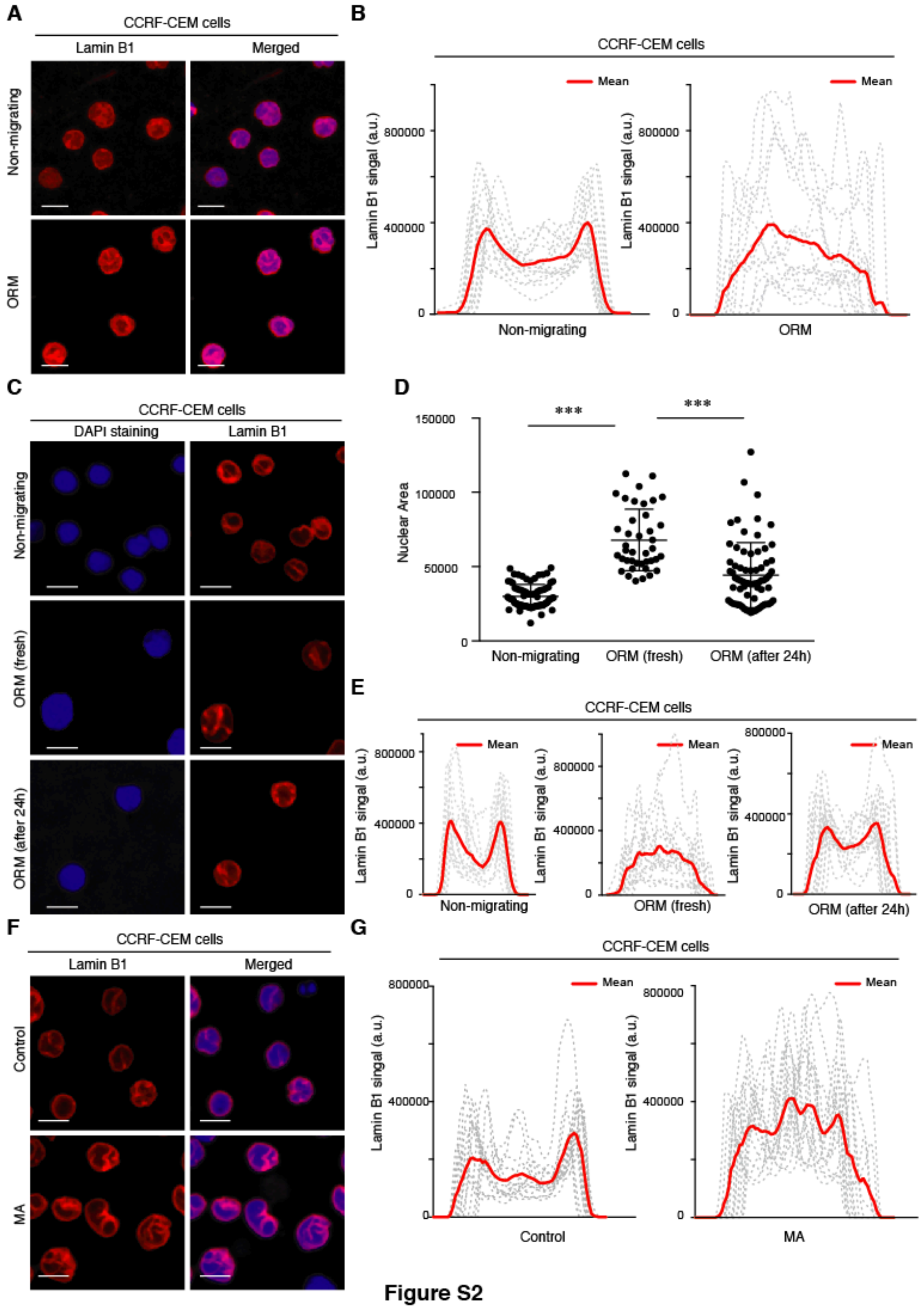


**Figure S2**

**Fig. S2. (A)** Non-migrating and ORM CCRF-CEM cells were seeded on poly-L-lysine-coated glasses and stained with DAPI (blue) and anti-lamin B1 antibody (red) for their analysis by confocal microscopy. Bar 10 μm. **(B)** Line plots show the signal profile of lamin B1 from 15 representative nuclei. Red line indicates the mean intensity of the profiles analyzed. **(C)** Non-migrating, fresh ORM, and ORM CCRF-CEM cells collected and cultured in suspension for an additional 24 h were seeded on poly-L-lysine-coated glasses and analyzed by confocal microscopy. Bar 10 μm. **(D)** Graph shows changes in the nuclear area of the cells from (C). Mean n=40-71 cells ± SD (2 replicates). **(E)** Line plots show the profiles based on lamin B1 intensity across 15 representative nuclei of the cells from (C). Red line indicates the mean intensity of the profiles analyzed. **(F)** Control and MA CCRF-CEM cells were seeded on poly-L-lysine-coated glasses and stained with DAPI (blue) and lamin B1 (red). Bar 10 μm. **(G)** Line plots show the signal profile of lamin B1 from 15 representative nuclei. Red line indicates the mean intensity of the profiles analyzed.

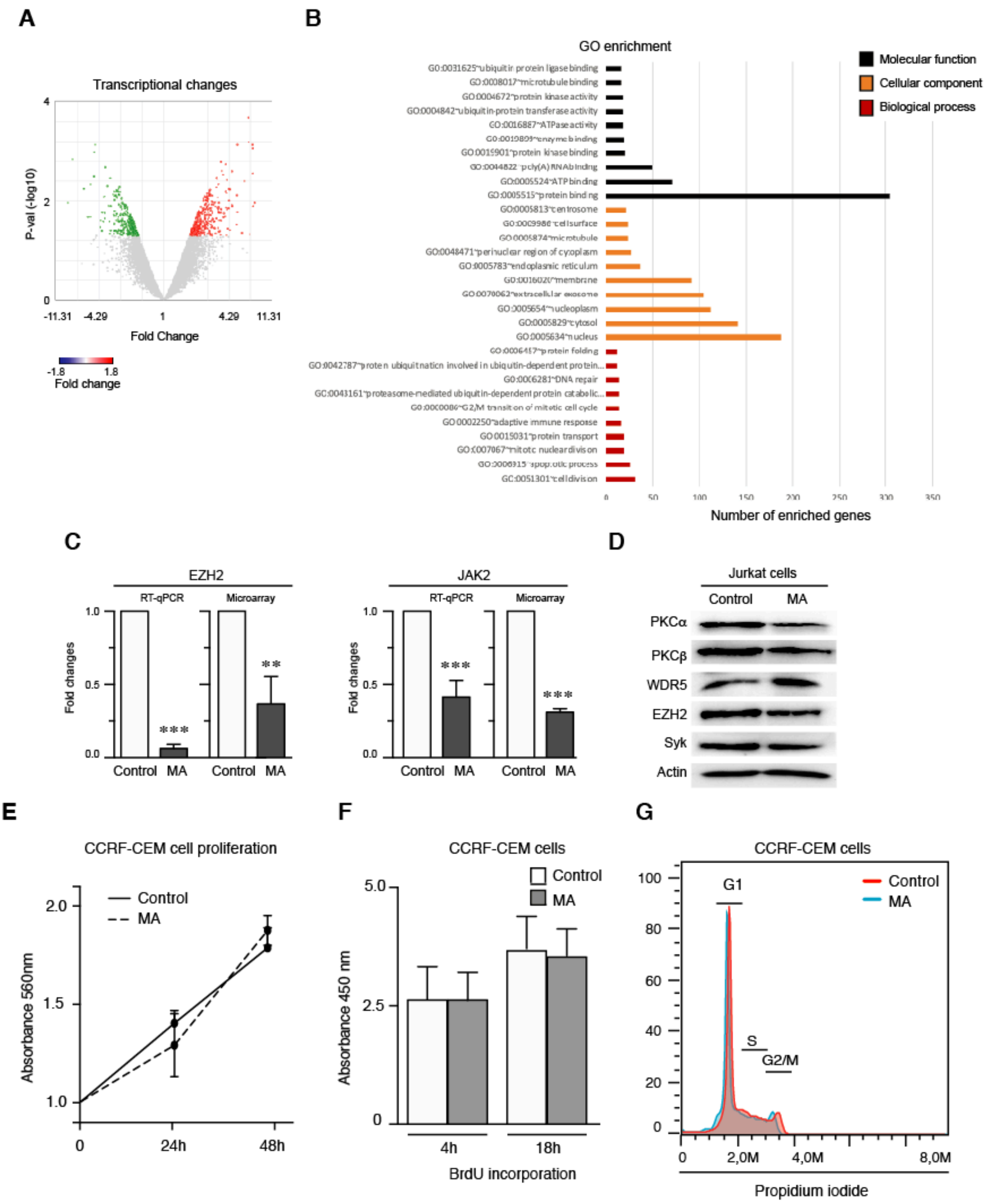


**Figure S3**

**Fig. S3. (A)** mRNA from control and MA Jurkat cells was isolated and the number of differentially expressed genes was analyzed by microarray. Volcano plot shows significant changes of gene transcripts in control and MA cells. **(B)** Graph shows the top gene ontology (GO) enrichment results from the microarray analysis. **(C)** Graph shows the validation of microarray by qPCR. The expression of EZH2 and JAK2 was detected in control and MA cells by qPCR (3 replicates) and the microarray data (2 replicates). Error bars indicate standard deviations. **(D)** Representative immunoblots for PKCα, PKCβ, syk, EZH2 and actin in the whole cell lysates of control and MA Jurkat cells. **(E)** Control (dark line) and MA (dashed line) CCRF-CEM cells were cultured at indicated times and cell proliferation was quantified by MTT assay. Mean n = 3 replicates ± SD. **(F)** Control and MA CCRF-CEM cells were incubated with BrdU for 4 and 18 h. Then, cells were fixed and BrdU incorporation was quantified. Mean n = 6 replicates ± SD. **(G)** Control (red) and MA (blue) CCRF-CEM cells were fixed, permeabilized and stained with propidium iodide. Then, cell cycle progression was analyzed by flow cytometry. Graph shows the G1, S and G2/M phases according to DNA content.

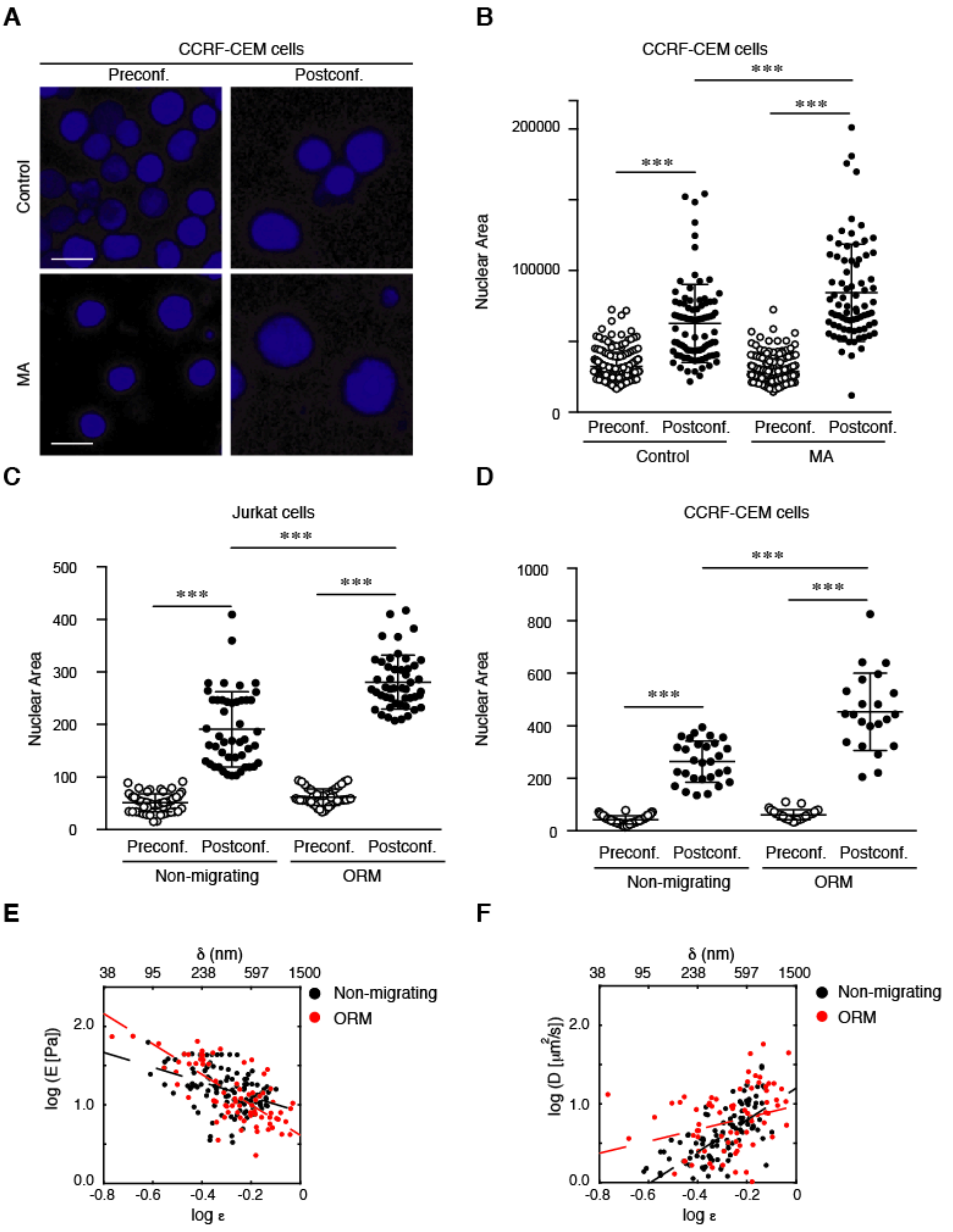


**Figure S4**

**Fig. S4. (A)** Isolated nuclei from control and MA CCRF-CEM cells were stained with DAPI and seeded on poly-L-lysine-coated coverslips. Nuclear area was determined before (Preconf.) and after (Postconf.) confinement. Bar 10 μm. **(B)** Graph shows the nuclear deformability upon mechanical compression. Mean n = 78-88 isolated nuclei ± SD (3 replicates). **(C)** Graph shows the nuclear deformability upon mechanical compression OF Isolated nuclei from non-migrating and ORM Jurkat cells. Nuclear area was determined before (Preconf.) and after (Postconf.) confinement. Bar 10 μm. Mean n = 47-59 isolated nuclei ± SD (3 replicates). **(D)** Graph shows the nuclear deformability upon mechanical compression OF Isolated nuclei from non-migrating and ORM CCRF-CEM cells. Nuclear area was determined before (Preconf.) and after (Postconf.) confinement. Bar 10 μm. Mean n = 22-63 isolated nuclei ± SD (3 replicates). **(E and F)** Variation of the stiffness (E) and the poroelastic diffusivity coefficient (D) in function of the nuclear indentation depth (shown in terms of strain, ε), both expressed as logarithmic values. Equivalent indentation depth in terms of distance, δ, is shown on top. Black and red dots represent values calculated from experimental data for non-migrated (n=109; indentation sites=12; nuclei=5) and migrated (n=79; indentation sites=10; nuclei=6) Jurkat cells, respectively, and linear fits are included as dashed lines.

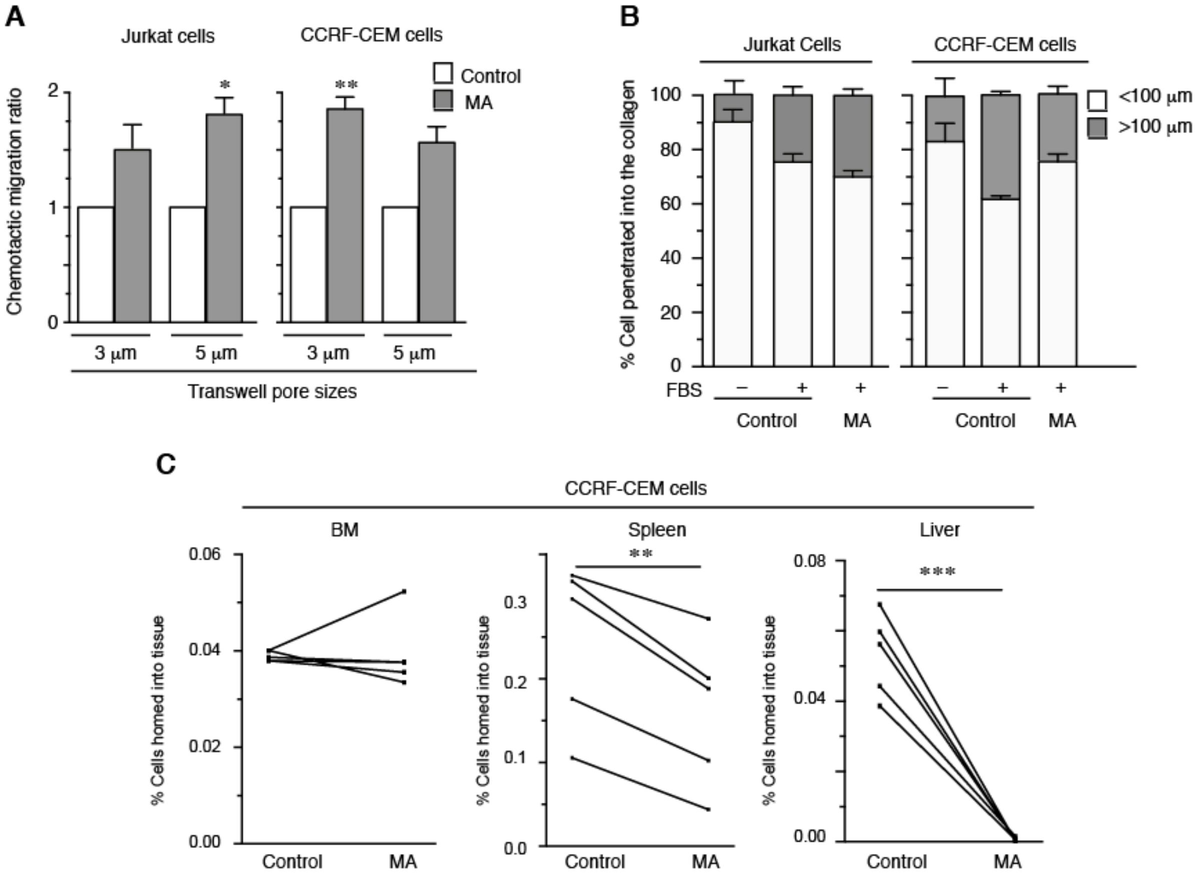


**Figure S5**

**Fig. S5. (A)** Control and MA Jurkat and CCRF-CEM cells were seeded on the top of Transwell chambers and allowed to migrate in response to serum (FBS). Cells were collected from the bottom chamber after 24 h and quantified. Mean n=3 replicates ± SD. **(B)** Control and MA Jurkat and CCRF-CEM cells were seeded on the top of a collagen matrix and allowed to penetrate into the collagen in response to serum (FBS, fetal bovine serum) for 24h. Cells were fixed, stained with propidium iodide and serial confocal sections were captured. Graph shows the percentage of cells invading deeper than 100 μm. Mean n=3 replicates ± SD. **(C)** Control (Cell Tracker Far Red+) and MA (CFSE+) CCRF-CEM cells were mixed 1:1 and injected into the tail vein of 5 NSG mice. After 24 h, mice were sacrificed, and labeled cells from spleen, liver and bone marrow were collected and determined by flow cytometry. Graph shows the percentage of control and the MA cells analyzed in each animal. Mean n = 5.

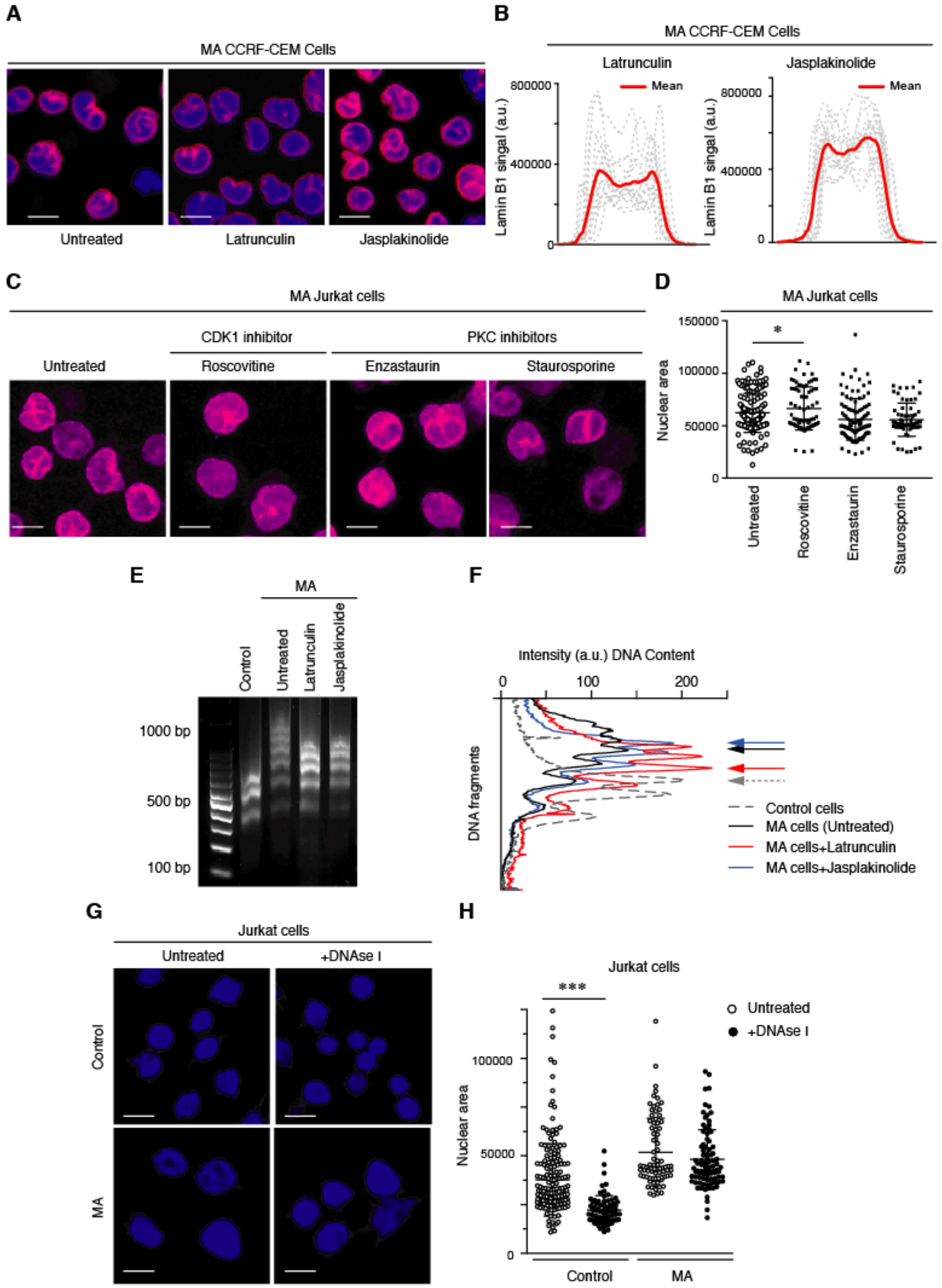


**Figure S6**

**Fig. S6. (A)** MA CCRF-CEM cells were cultured in the presence or absence of latrunculin B (1 µg/mL) and japlakinolide (1 µg/mL) for 1 h at 37ºC. Then, cells were seeded on poly-L-lysine coated coverslips, fixed, permeabilized and stained with DAPI (blue) and anti-laminB1 (red) antibody. Bar 10 μm. **(B)** Line plots show the signal profile of lamin B1 from 15 representative nuclei. Red line indicates the mean intensity of the profiles analyzed. **(C)** Control and MA Jurkat cells were cultured in the presence or absence of roscovitine (15 μM, Cdk1 inhibitor), enzastaurin and staurosporine (50 nM, 2 nM, respectively, PKC inhibitors), for 30 min. Then, cells were fixed, permeabilized and stained with DAPI and anti-laminB1 antibody. Bar 10 μm. **(D)** Graph shows changes in the nuclear area of MA cells upon treatments in (C). Mean n=65-186 cells ± SD (3 replicates). **(E)** Control and MA Jurkat cells were incubated with indicated treatments, collected, and their DNA was digested with DNAse for 8 min. Then, DNA fragments were resolved in an agarose gel. **(F)** Graph shows the nucleosomal releasing profile from control (dashed line) and MA Jurkat cells as in (E). Arrows indicate the maxima DNA peaks in each cell population. **(G)** Control and MA Jurkat cells were collected, and their DNA was digested with DNAse for 20 min. Then cells were stained with DAPI (blue), fixed, and analyzed by confocal microscopy. Bar 10 μm. **(H)** Graph shows changes in the nuclear area of control and MA Jurkat cells as in (G). Mean n=72-185 cells ± SD.

#### SUPPLEMENTARY TABLES

**Supplementary Table S1**. Transcriptional changes of control and MA Jurkat cells by microarray analysis. |Fold Change| > 1.8 and P-value <0.05. n=2.

**Supplementary Table S2**. The table shows the terms from the microarray analysis with the highest significance according to the adjusted FDR p-value, ordered by pathway.

#### SUPPLEMENTARY MOVIES

**Supplementary Movies S1-S4**. 3D reconstructions from confocal sections of representative DAPI-stained nuclei from non-migrating (S1), ORM (S2), control (S3) and MA (S4) Jurkat cells.
